# Carotid intima-media thickness in UK Biobank: Identification of novel genome-wide loci, sex-specific effects and genetic correlations with obesity and glucometabolic traits

**DOI:** 10.1101/718684

**Authors:** Rona J. Strawbridge, Joey Ward, Mark E.S. Bailey, Breda Cullen, Amy Ferguson, Nicholas Graham, Keira J.A. Johnston, Laura M. Lyall, Robert Pearsall, Jill Pell, Richard J Shaw, Rachana Tank, Donald M. Lyall, Daniel J. Smith

## Abstract

**Objectives:** Atherosclerosis is the underlying cause of most cardiovascular disease, but mechanisms underlying atherosclerosis are incompletely understood. Ultra-sound measurement of the carotid artery intima-media thickness (cIMT) can be used to measure vascular remodelling, which is indicative of atherosclerosis. Genome-wide association studies have identified a number of genetic loci associated with cIMT, but heterogeneity of measurements collected by many small cohorts have been a major limitation in these efforts. Here we conducted genome-wide association analyses in UK Biobank (N=22,179), the largest single study with consistent cIMT measurements.

**Approach and results:** We used BOLT-LMM to run linear regression of cIMT in UK Biobank, adjusted for age, sex, genotyping platform and population structure. In white British participants, we identified 4 novel loci associated with cIMT and replicated most previously reported loci. In the first sex-specific analyses of cIMT, we identified a female-specific locus on Chromosome 5, associated with cIMT in women only and highlight *VCAN* as a good candidate gene at this locus. Genetic correlations with body-mass index and glucometabolic traits were also observed.

**Conclusion:** These findings replicate previously reported associations, highlight novel biology and provide new directions for investigating the sex differences observed in cardiovascular disease presentation and progression.

## Introduction

Atherosclerosis is the underlying cause of the majority of cardiovascular events (CVE) and is characterised by vascular remodelling, incorporation of lipids into the vessel wall and subsequent inflammation ^1, 2^. Atherosclerosis is a systemic process which precedes clinical presentation of cardiovascular events such as stroke by decades. Indeed, evidence of vascular remodelling indicative of atherosclerosis has been observed as early as in adolescent age groups ^3^.

Atherosclerosis can be non-invasively assessed by ultrasound measurement of the carotid artery vessel wall, specifically the intima-media thickness (cIMT). In some cases, cIMT assessment is used for monitoring after CVE such as stoke, but could also be useful for screening individuals at high risk of CVE. Currently use is limited as it requires specialist equipment and training, and high-quality data analysis is laborious. Measurement of cIMT has been performed for research purposes, predominantly in cohorts recruited for the study of cardiovascular disease. Whilst undeniably useful, the use of clinical cohorts does not cover the whole spectrum of atherosclerotic burden in the population.

Genetic analyses of clinical cohorts have begun to identify single nucleotide polymorphisms (SNPs) associated with increased cIMT ^4–7^, which paves the way for better understanding of processes leading to cardiovascular events. A limitation for these studies (N∼68,000) has been heterogeneity in recruitment and ultrasound methodology, which could lead to failure to detect some true genetic effects. In this respect, UK Biobank provides an unprecedented opportunity to analyse IMT measurements in a very large cohort (N∼22,000) with consistent recruitment and standardised cIMT measurements, analysis and quality control.

We therefore set out primarily to identify genetic variants associated with cIMT in a large general population cohort. A secondary aim was to investigate the possibility of sex-specific genetic effects on IMT. Here we report the replication of previously reported associations, highlight novel biology and provide new directions for investigating the sex differences observed in cardiovascular disease presentation and progression.

## Methods

### Study population

The UK Biobank study (UKB) has been described in detail previously ^8, 9^. In brief, UKB recruited ∼500,000 participants from the UK between 2006 and 2010. Participants attended one of the 22 recruitment centres across the UK where they provided a blood sample for DNA extraction and biomarker analysis, and completed questionnaires covering a wide range of medical, social and lifestyle information. All participants provided informed consent and the study was conducted in accordance with the Helsinki Declaration. Generic approval was granted by the NHS National Research Ethics Service (approval letter dated 13 May 2016, Ref 16/NW/0274) and the study conducted under UK Biobank projects #7155 (PI Jill Pell) and #6553 (PI Daniel Smith).

### Phenotyping

Starting in 2014, participants were invited to participate in a follow-up and imaging assessment, including ultrasound imaging of the carotid arteries. cIMT phenotyping began in 2015, in a pilot phase, where 2,272 individuals were at imaged at 18 centres (with 8 centres accounting for 98% of the sample) with extensive manual quality control being conducted. Subsequently, manual quality control was deemed unnecessary and all centres began recruiting and recording automated measurements (with 10 centres accounting for 93% of the sample). Details of the protocol are available at https://biobank.ctsu.ox.ac.uk/crystal/label.cgi?id=101. Inbrief, ultrasound measurements were recorded at 2 angles on each of the left and right carotid artery with automated software recording images and measurements of mean and maximum intima-media (UKB data fields 22670-22681). Recruitment for imaging is ongoing, but to date (2019), 25,769 individuals have ultrasound measurements of the cIMT. The average of 4 measures (2 for each of the left and right carotid arteries) was calculated for the mean (IMTmean) and for maximum IMT (IMTmax), the largest of the 4 measures was used. Where more than one value was missing due to poor quality of the image, the participant was excluded from analyses. Values were expressed in mm and natural log transformed for normality prior to analysis.

In the follow-up assessment, anthropometric measures, lifestyle variables, medication and disease history was again recorded including: age, weight, waist circumference, hip circumference, waist:hip ratio (WHR), body-mass index (BMI), systolic and diastolic blood pressure (SBP and DBP respectively), hypertension (defined as SBP≥140mmHg and/or DBP≥90mmHg and/or anti-hypertensive medication), probable type 2 diabetes (T2D, coding as per Eastwood et al ^10^), ischemic heart disease (ISH, defined as heart attack or angina). Corrected SBP and DBP, reflecting probable untreated levels, were calculated as per Ehret et al ^11^. These contemporary values were used in the analysis of cIMT.

### Genotyping

DNA was extracted from blood samples provided by participants, using standard protocols. Details of the UKB genotyping and imputation procedures have been described previously ^12, 13^. Briefly, the full genetic data release (March 2018) was used for this study. Genotyping, pre-imputation quality control, imputation and post-imputation quality control were conducted centrally by UKB, according to standard procedures.

### Statistical analyses

Descriptive statistics and Spearmans rank correlations were conducted using Stata. Only individuals of white British ancestry were included in the GWAS to maximise homogeneity. BOLT-LMM was used to conduct genetic association analyses, to calculate heritability estimates and estimates of λGC. IMTmean and IMTmax values were natural logarithm-transformed for normality and genetic association analyses were conducted, adjusted for age, sex and genotyping array (primary analysis) or age and genotyping array (secondary analyses). SNPs were excluded if minor allele frequency <0.01, Hardy-Weinberg equilibrium p<1×10^-6^ or imputation score <0.3. Genome-wide significance was set at p<5×10^-8^, with suggestive evidence of association being set at p<1×10^-5^. After quality control there were 22,179 participants with IMT and genetic data for analysis. Genetic association results were visualised using FUMA ^14^ and LocusZoom ^15^.

### Linkage disequilibrium and genetic correlations

Linkage disequilibrium (LD) between analysed SNPs in each GWAS-significant locus was calculated and visualised in a random subset of 10000 white British individuals (or 5000 individuals where the locus is computationally too large with 10000 individuals) included in the cIMT subset, using Haploview (default settings) ^16^.

Genetic correlations between IMTmean and IMTmax and relevant cardiometabolic traits were calculated, using previously published summary statistics and LD score regression ^17^. IMT summary statistics were provided by the CHARGE consortium (http://www.chargeconsortium.com/). Data on glycaemic traits were contributed by MAGIC investigators and were downloaded from www.magicinvestigators.org. Type 2diabetes (T2D) data were contributed by the DIAGRAM consortium (http://diagram-consortium.org/downloads.html). Summary statistics for lipid traits were downloaded from the Global Lipids Genetics Consortium website (http://lipidgenetics.org/). Data on anthropometric traits were downloaded from the GIANT consortium website (http://portals.broadinstitute.org/collaboration/giant/index.php/Main_Page). Coronary artery disease data were downloaded from the CARDIoGRAMplusC4D consortium (http://www.cardiogramplusc4d.org/). Blood pressure data were provided by the International Consortium for Blood Pressure genetics (https://www.ncbi.nlm.nih.gov/projects/gap/cgibin/study.cgi?study_id=phs000585.v1.p1).

### Data-mining

The GWAS catalogue (https://www.ebi.ac.uk/gwas/, accessed 20190717) was used to identify previously reported associations with significant loci. All SNPs with at least suggestive evidence of association (p<1×10^-5^) within significantly associated loci were assessed for deleterious effect using the Ensemble variant effect predictor (VEP) ^18^. GWAS-significant SNPs and those predicted by VEP to have at least moderate impact were assessed for effects on genotype expression patterns of nearby genes using the Genotype Tissue Expression project (GTEx) ^19^. Functions of highlighted genes were explored using the NCBI Gene platform (https://www.ncbi.nlm.nih.gov/gene/, accessed 20190717) and literature identified using NCBI PubMed (https://www.ncbi.nlm.nih.gov/pubmed/, accessed 20190717).

## Results

Demographic characteristics of the UKB cIMT subset are presented in Table 1. The UKB cIMT subset consist of 48.3% men. Women were slightly younger (average 54.6 years versus 56.0 years) and were healthier (average BMI 26.1 kg/m2 and SBP 132 mmHg, % with hypertension 37.0%) and smoked less (% smokers 30.4) compared to men (average 56.0 years, BMI 27.1 kg/m2, 140 mmHg, % hypertension 54.0, smokers 36.0). Values for both IMTmean and IMTmax were lower in women than in men. Measurements of IMTmean and IMTmax were highly correlated (Spearmans Rho=0.865, p<0.0001 (full cIMT subset), Rho=0.868 p<0.0001 (men) and Rho=0.852 p<0.0001 (women)). IMTmean and IMTmax were significantly and positively associated with classical risk factors including increasing age, obesity, and blood pressure (Table 2).

**Table 1:** Demographics of the UKB cohort with cIMT measures

|  | Men | Women | all |
| --- | --- | --- | --- |
| N (%) | 10708 (48.3) | 11471 (51.7) | 22179 |
| Age (years) | 63.9 (7.6) | 62.4 (7.3) | 63.1 (7.5) |
| Weight (Kg) | 83.8 (13.2) | 69.2 (13.9) | 76.2 (15.0) |
| Waist (cm) | 94.0 (10.4) | 82.6 (11.5) | 88.1 (12.4) |
| Hip (cm) | 101.2 (7.2) | 101.4 (9.8) | 101.3 (8.6) |
| WHR | 0.93 (0.06) | 0.81 (0.07) | 0.87 (0.09) |
| BMI (kg/m<sup>2</sup>) | 27.0 (3.8) | 26.1 (4.7) | 26.5 (4.3) |
| SBP (mmHg) | 141 (17) | 134 (18) | 137 (18) |
| DBP (mmHg) | 80 (10) | 77 (10) | 79 (10) |
| Corrected SBP (mmHg)* | 145 (19) | 137 (20) | 141 (10) |
| Corrected DBP (mmHg)* | 83 (11) | 79 (11) | 81 (11) |
| IMTmean (mm) | 0.706 (0.135) | 0.658 (0.109) | 0.682 (0.125) |
| IMTmax(mm) | 0.952 (0.216) | 0.875 (1.86) | 0.912 (0.204) |
| N hypertension | 7320 (69.0) | 6121 (53.8) | 13441 (61.2) |
| N on BP medication | 3026 (28.4) | 1973 (17.3) | 4999 (22.7) |
| N on lipid lowering medication | 3255 (34.5) | 1681 (17.7) | 3255 (34.5) |
| N probable t2d | 614 (5.7) | 325 (2.8) | 939 (4.2) |
| N ISH | 275 (2.6) | 97 (0.9) | 372 (1.7) |
| n current smoking | 444 (4.2) | 356 (3.1) | 800 (3.6) |

Where: \* adjusted for blood pressure medication as per Ehret et al, T2D, type 2 diabetes; ISH, ischemic heart disease

**Table 2:**
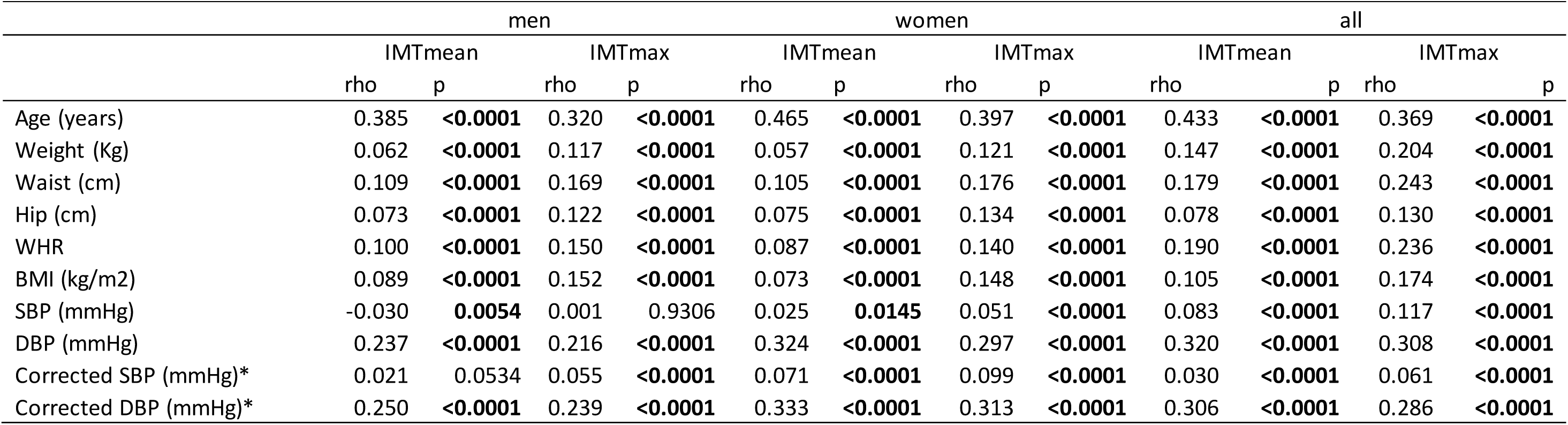
Correlations between cIMT and classical cvd risk factors

### Primary analysis: SNPs associated with IMT_mean_ and IMT_max_

Manhattan and QQ plots of the GWAS results are presented in Figure 1. There was some evidence of inflation for both IMTmean and IMTmax (λ_GC_=1.15 and 1.10, respectively), however this is likely due to polygenicity rather than population structure (LDSR intercept (standard error) = 1.03 (0.03) for both phenotypes).

GWAS-significant evidence of association with IMTmean was observed for 176 SNPs in 8 loci (Figure 1A) and 76 SNPs in 3 loci demonstrated GWAS-significant evidence of association with IMTmax (Figure 1B). The lead SNP for each locus is provided in Table 3. As BOLT-LMM includes neighbouring SNPs in the model (essentially conditioning on other SNPs in the region), each locus reported here is independent and contains only a single signal. Effect sizes of all lead SNPs were comparable with those previously reported (Beta 0.0091-0.441) ^6^.

Of the 8 loci significantly associated with IMTmean identified here, four are novel (Chr7, lead SNP rs342988; Chr8 (124.6Mb), rs34557926; Chr11, rs2019090; Chr19 (41.3Mb), rs111689747; Figure 2A-D) and 4 have previously been reported (Figure 2E-H).

**Figure 1:**
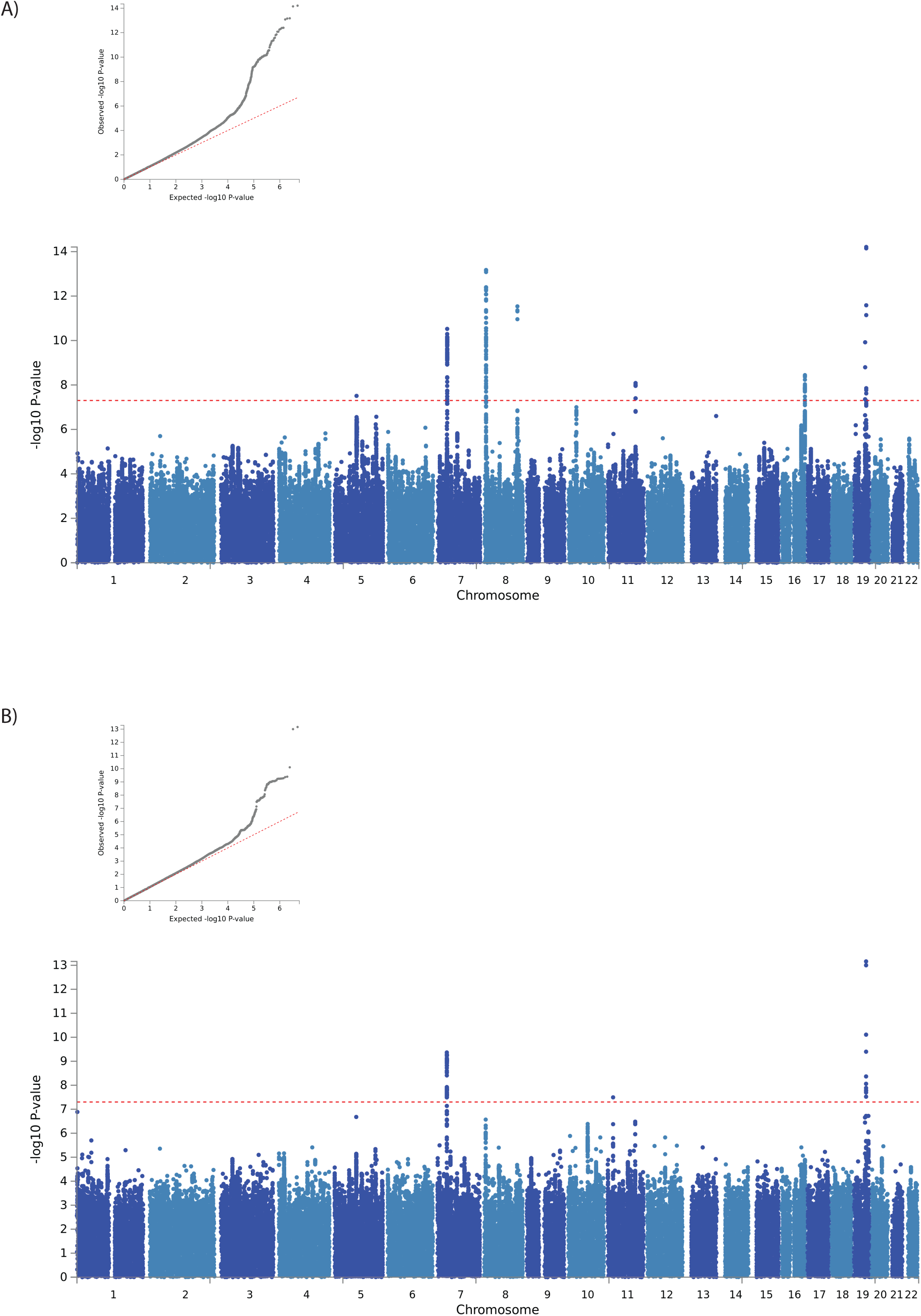
QQ and Manhattan plots of results of the GWAS of A) IMTmean andB) IMTmax.

**Figure 2:**
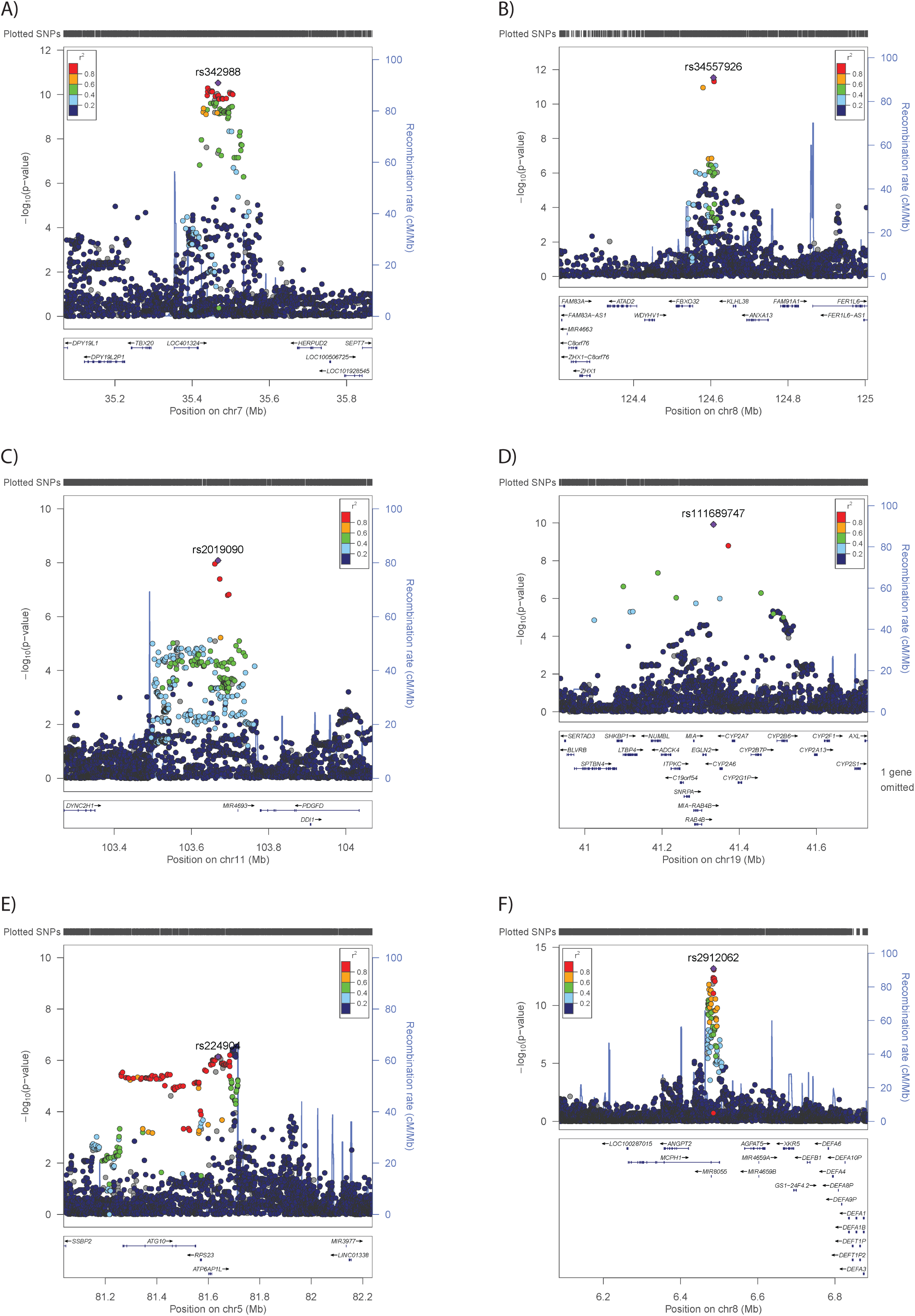
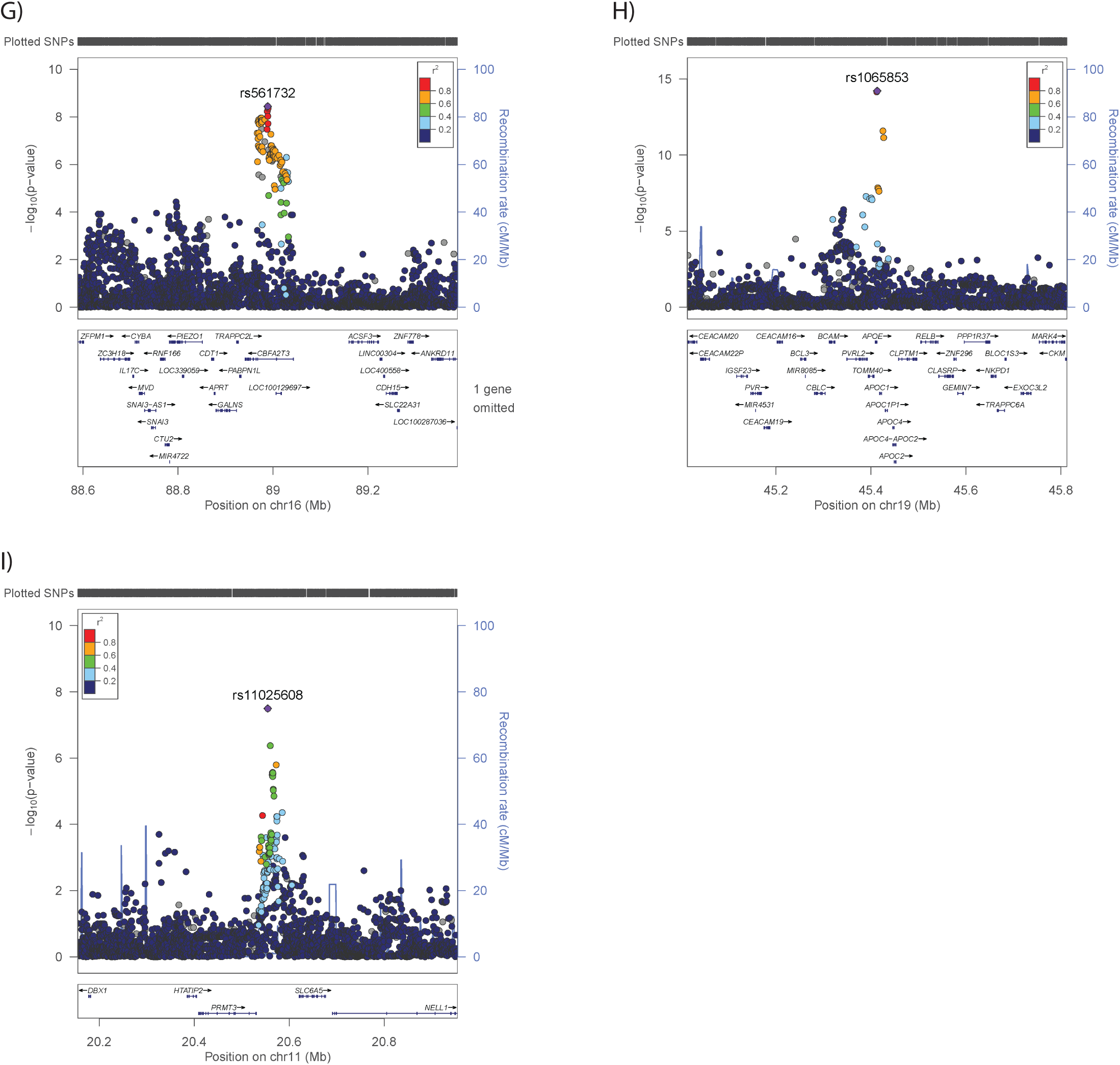
Regional plots for novel IMTmean-associated loci on A) Chr7 (rs342988), B) Chr8 (rs34557926), C) Chr11 (rs2019090), D) Chr19 (rs111689747) and known IMTmean-associated loci on E) Chr5 (rs224904, instead of lead SNP rs758080886), F), Chr8 (rs2912062), G) Chr16 (rs561732), H) Chr19 (rs1065853) as well as the novel locus for IMTmax on Chr11 (rs11025608). The lead SNP is indicated by a purple diamond. LD (r2) between other SNPs and the lead SNP is indicated by colour. Grey indicates LD is not known.

**Table 3:** lead SNPs in loci assocaited with cIMT

| Analysis | SNP | CHR | locus* | Genes under the locus | A1 | A0 | A1<br>freq | beta | se | chi-2 | P | GWAS<br>SNPs<br>in<br>locus |
| --- | --- | --- | --- | --- | --- | --- | --- | --- | --- | --- | --- | --- |
| IMTmean<br>sex<br>combined | rs758080886 | 5 |  | *ATG10, RPS23, ATP6AP1L* | CA | C | 0.15 | 0.013 | 0.002 | 30.6 | 3.10E-08 | 1 |
|  | rs342988<sup>a, d</sup> | 7 | 35423859-35528567 | *LOC401324* | C | T | 0.28 | 0.011 | 0.002 | 44.2 | 3.00E-11 | 78 |
|  | rs2912062<sup>b</sup> | 8 | 6470650-6494850 | *MCPH1, MIR8055* | C | G | 0.71 | 0.012 | 0.002 | 56.2 | 6.70E-14 | 56 |
|  | rs34557926<sup>d</sup> | 8 | 124579985-124608614 | *FBXO23, KLHL38, ANAXA13* | C | T | 0.64 | -<br>0.011 | 0.002 | 48.8 | 2.90E-12 | 4 |
|  | rs2019090<sup>d</sup> | 11 | 103660567-103673294 | *MIR4693* | A | T | 0.29 | 0.009 | 0.002 | 33.2 | 8.20E-09 | 5 |
|  | rs561732 | 16 | 88966667-88989862 | *CBFA2T3, LOC100129697,*<br>*TRAPPC2L* | T | C | 0.65 | 0.009 | 0.002 | 34.8 | 3.60E-09 | 22 |
|  | rs111689747<sup>d</sup> | 19 | 41188985-41371458 | *CYP2A6, CYP2A7,*<br>*CYP2G1PCYP2B7P, CYP2B6,*<br>*SNRPA, MIA, EGLN2, RAB4B,*<br>*MIA-RAB4B, RAB4B-EGLN2,*<br>*C19ORF54, ITPKC, ADCK4,*<br>*NUMBL, LTBP4, SHKBP1,*<br>*SPTBN4* | G | A | 0.99 | -<br>0.044 | 0.007 | 41.5 | 1.20E-10 | 3 |
|  | rs1065853<sup>c</sup> | 19 | 45412079-45426792 | *BCAM, PVRL2, TOMM40, APOE,*<br>*APOC1, APOC1P1, APOC4-*<br>*APOA2, APOC2, CLPTM1,*<br>*APOC4, RELB* | G | T | 0.92 | 0.021 | 0.003 | 60.8 | 6.20E-15 | 7 |
| IMTmax<br>sex<br>combined | rs11762074<sup>a</sup> | 7 | 35428513-35505204 | *LOC401324* | T | C | 0.77 | -<br>0.014 | 0.002 | 39 | 4.30E-10 | 63 |
|  | rs11025608<sup>d</sup> | 11 |  | *PRMT3, SLC6A5* | G | T | 0.44 | -<br>0.010 | 0.002 | 30.6 | 3.20E-08 | 1 |
|  | rs1065853<sup>c</sup> | 19 | 45389596-45426792 | BCAM, PVRL2, TOMM40, APOE, APOC1, APOC1P1, APOC4-APOA2, APOC2, CLPTM1, APOC4, RELB | G | T | 0.92 | 0.026 | 0.003 | 56.1 | 6.90E-14 | 11 |
| IMTmean Men | rs35099106<sup>a</sup> | 7 | 35428513-35505204 | LOC401324 | C | CT | 0.24 | 0.016 | 0.003 | 34.1 | 5.20E-09 | 38 |
|  | rs2912063<sup>b</sup> | 8 | 6470650-6490544 | MCPH1, MIR8055 | A | G | 0.71 | 0.017 | 0.003 | 44.5 | 2.60E-11 | 24 |
|  | rs1065853<sup>c</sup> | 19 | 45412079-45413233 | BCAM, PVRL2, TOMM40, APOE, APOC1, APOC1P1, APOC4-APOA2, APOC2, CLPTM1, APOC4, RELB | G | T | 0.92 | 0.027 | 0.004 | 39.1 | 4.00E-10 | 2 |
| IMTmean Women | rs309563<sup>d</sup> | 5 | 82870033-82933913 | VCAN, HAPLN1 | A | T | 0.66 | -0.01 | 0.002 | 37.6 | 8.90E-10 | 37 |

Where: \*, region bordered by gwas-significant SNPs; a, LD between rs342988, rs11762074 and rs35099106 suggests that these SNPs represent 1 signal; b, LD between rs2912062 and rs2912063 indicate that these SNPs are proxies; c, consistent lead SNP convincingly demonstrates that the same locus is being identified; d, novel locus.

In the previously reported loci, on Chr5 there is only one SNP, rs758080886, which reaches GWAS-significant evidence for association with IMTmean which is located 3417b from the previously reported lead SNP rs224904 ^6^. rs758080886 is not available in the reference panel used by LocusZoom ^15^ and is therefore not plotted, hence this region is represented by the previously reported SNP, rs224904. The LD between rs75808088 and rs224904 is moderate (r2=0.44) therefore rs224904 is not a good proxy, however this is the same for all SNPs in the locus (SFigure A). On Chr8, rs2912062 is 738b from the previously reported lead, rs2912063, and in high LD (r2=0.98), demonstrating that these SNPs represent the same signal. On Chr16, the lead SNP, rs561732, is 22.3kb from the previously reported lead, rs844396, with moderate LD (r2=0.66). The Chr19 lead here, rs1065853 is 1.15kb from the reported rs7412, but with almost complete LD (r2=0.99). The novel Chr19 locus is ∼4Mb from this signal, and whilst long-range LD is possible, the calculated LD between rs111689747 and rs1065853 or rs7412 does not support this possibility (r2=0).

There were three GW-significant loci for IMTmax. Of these, two overlap with those for IMTmean: the lead SNPs for IMTmean and IMTmax on Chr7 (rs342988 and rs11762074 respectively) are 26.9kb apart with LD of r2=0.77. For Chr19, the locus not only overlaps, but the lead SNP is the same for both traits. The Chr11 for IMTmax (20.5Mb) is distinct from that for IMTmean (103.6Mb) (Figure 2I).

When considering SNPs significantly associated with IMTmean, the direction of effects on IMTmax are consistent with those for IMTmean, and the magnitude is similar (STable 1). The same is true for the Chr11 IMTmax SNP, rs11025608, which shows a similar effect size and direction in IMTmean (beta −0.008, se 0.002, p=1.6×10^-6^). Therefore further analyses focused on IMTmean, for more robust comparisons with previous studies.

### UKB IMT GWAS and the CHARGE consortium IMT GWAS meta-analysis

The largest GWAS meta-analysis of IMTmean previously published ^6^ included 68,962 individuals from 30 studies with a variety of recruitment strategies, inclusion criteria and measurements of cIMT. The range of average ages was 37.7-78.8 years and average IMTmean 0.50-1.13mm. The single largest study within the meta-analysis included 8,663 individuals (approximately evenly split between men and women. That meta-analysis reported 11 robustly associated loci^6^, of which nine lead SNPs were available in UKB (Stable 2). Seven of these demonstrated significant (p<0.05) associations with IMTmean, with consistent effect directions when compared to the previous report ^6^ (Stable 2). One SNP demonstrated a non-significant association and one demonstrated a significant association but inconsistent direction (Stable 2).

Conversely, of the lead SNPs reported here for UKB, 10 were available in the CHARGE meta-analysis (Stable 3). Of these, nine were significant (p<0.05) with consistent effect directions to those reported in UKB. One lead SNP was not significant. Effects sizes were generally 2-3-fold larger in UKB than in the CHARGE meta-analysis.

### Secondary analyses: sex-specific genetic effects on IMT

Sex-specific analyses suggest that the genetic variants associated with IMTmean in men and women is distinctly different (Figure 3): In men-only analyses (Figure 3A), three GWAS-significant loci were identified on Chr7 (rs35099106), Chr8 (6.4Mb, rs2912063) and Chr19 (rs1065853), all of which overlap with those identified in the primary (sex-combined) analysis (Table 3 and Figure 4). The Chr7 lead SNP is in strong LD with that for IMTmax (rs11762074, r2=0.97) but only moderate LD with the sex-combined IMTmean lead SNP (rs342988, r2=0.64. The lead SNPs on Chr8 and 19 are consistent with either the primary analysis or with previously reported lead SNPs.

**Figure 3:**
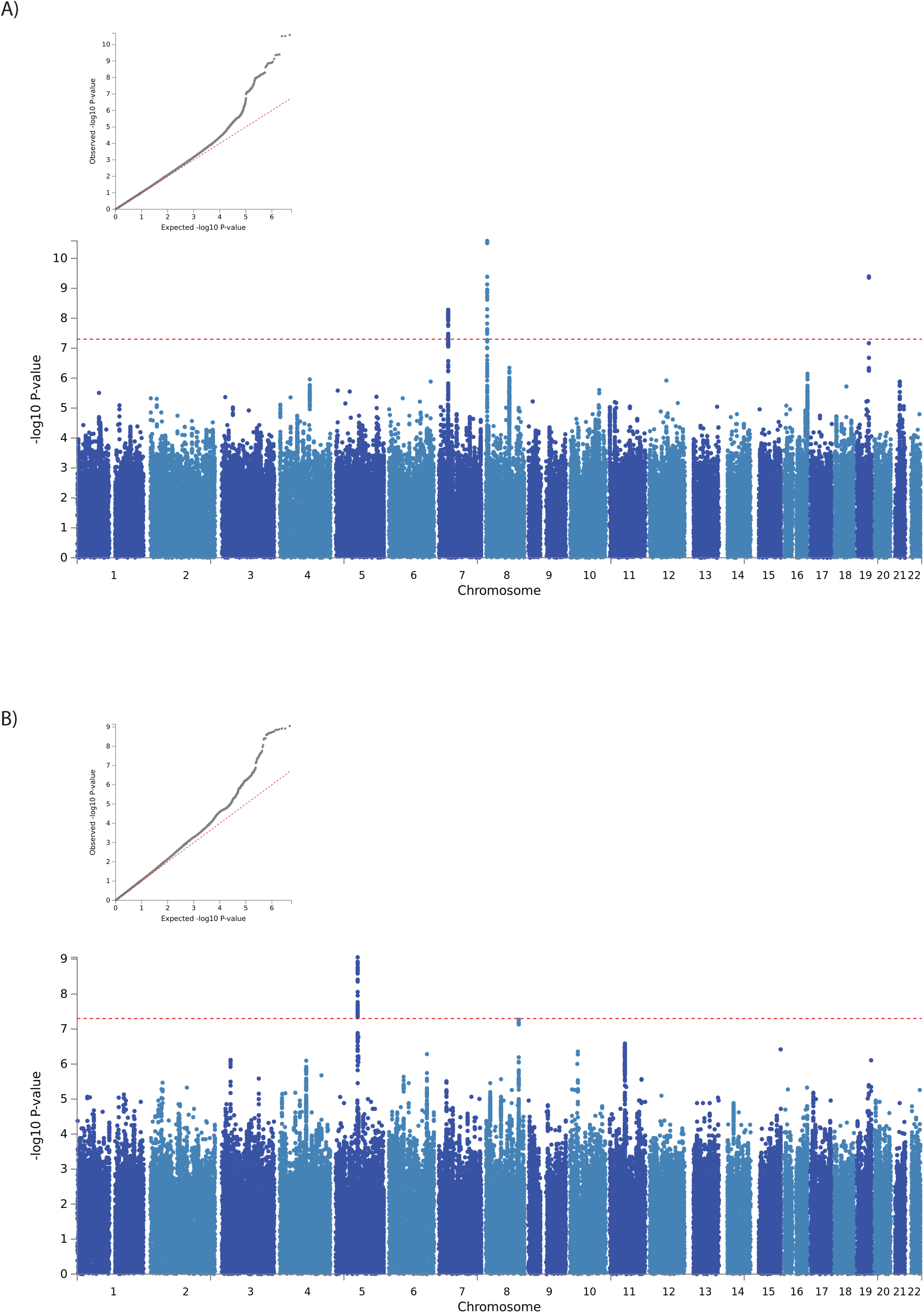
QQ and Manhattan plots of results of the GWAS of IMTmean in A) men only and B) women only.

**Figure 4:**
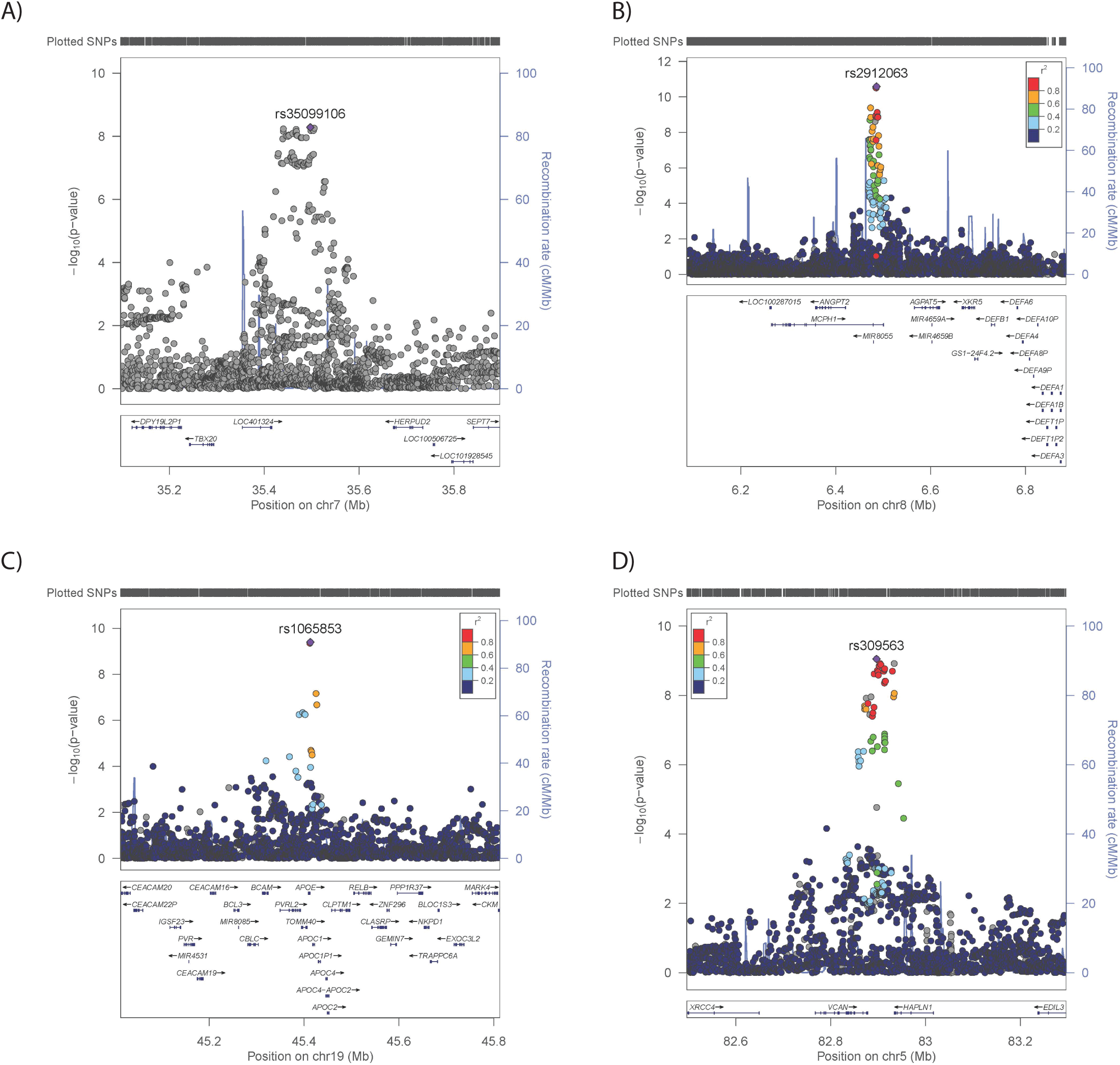
Regional plots for sex-specific IMTmean-associated loci. Men-only loci on A) Chr7 (rs35099106), B) Chr8 (rs2912063), C) Chr19 (rs1065853) and the women-only locus on D) Chr5 (rs309563). The lead SNP is indicated by a purple diamond. LD (r2) between other SNPs and the lead SNP is indicated by colour. Grey indicates LD is not known.

In analysis of women (Figure 3B), only a locus on Chr5 was GWAS-significant (Table 3 and Figure 4). The lead SNP for this locus, rs309563, is ∼1.2Mb from, and demonstrated no LD with, the lead sex-combined SNP for IMTmean (r2=0) or the previously reported lead SNP for this locus (rs224904, r2=0, SFigure A), suggesting that it is a separate locus.

Effect directions for SNPs with suggestive evidence of association were consistent between men and women, however in men, the SNPs associated with cIMT in women demonstrated 3-6 fold smaller (non-significant) effect sizes (STable 4). In contrast, for the SNPs associated with cIMT in men, effect sizes were halved in women, with at least nominal significance being observed for most SNPs, and suggested significance noted for the Chr19/APOE locus.

### Genetic correlations with cardiovascular phenotypes and risk factors

UKB IMTmean and IMTmax GWAS demonstrated significant genetic correlations with the CHARGE GWAS IMTmean meta-analyses (rg =0.77-0.82, Table 4) ^4, 6^. As the individuals included in the IMTmean and IMTmax analyses overlap completely was not possible to compare these analyses. Significant positive genetic correlations were observed between IMTmean and total obesity (BMI), type 2 diabetes (T2D), fasting glucose and insulin (Table 4). For IMTmax, positive correlations with total and central obesity (BMI and WHRadjBMI), T2D, fasting glucose and rheumatoid arthritis were observed. It was surprising to note that a significant positive association with high density lipoprotein (HDL) and a significant negative association with fasting insulin was observed. The significance of these findings is unclear.

**Table 4:**
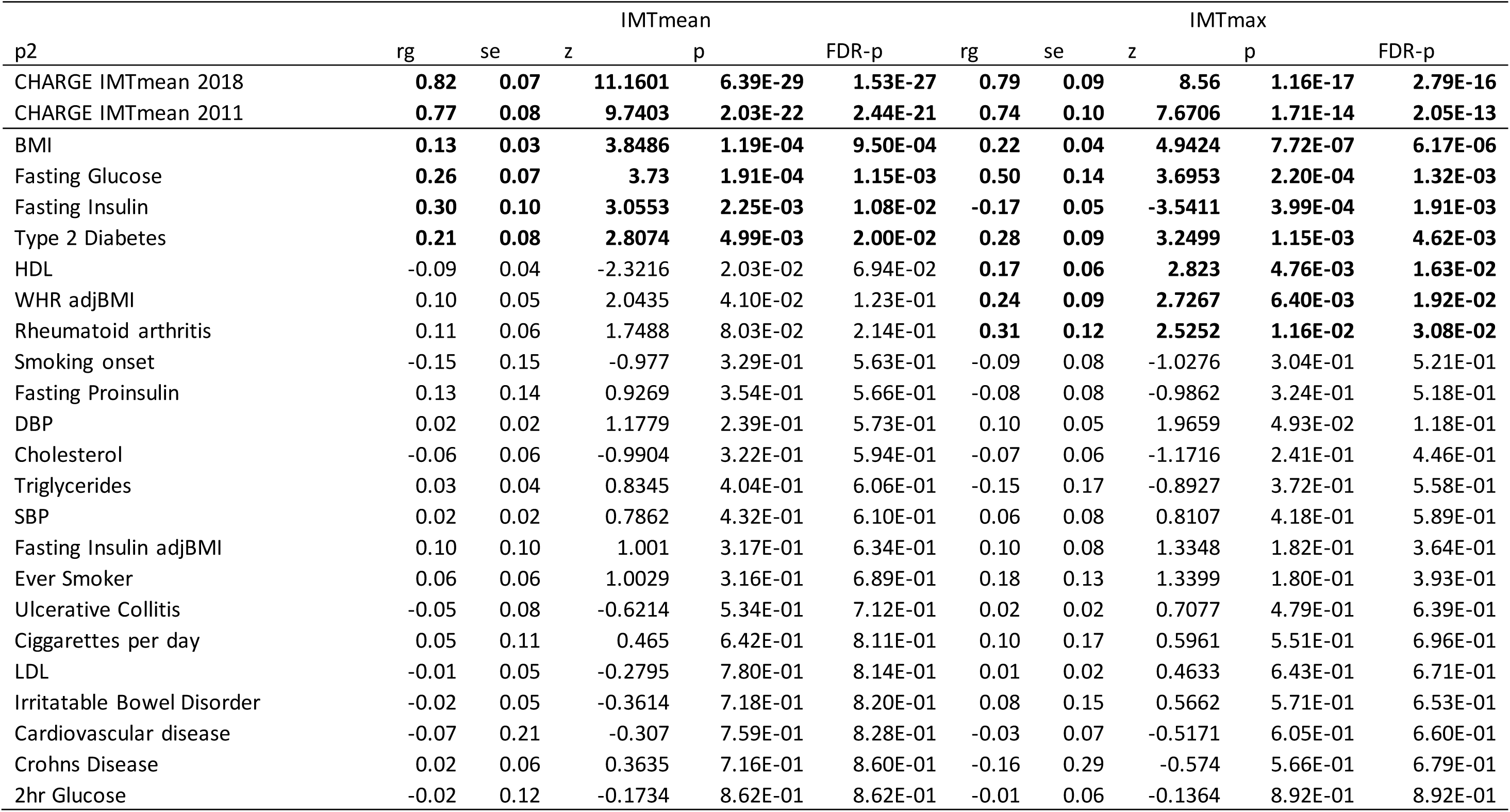
Genetic correlations between IMTmean, IMTmax and cardiometabolic traits and risk factors

| p2 | IMTmean |  |  |  |  | IMTmax |  |  |  |  |
| --- | --- | --- | --- | --- | --- | --- | --- | --- | --- | --- |
|  | rg | se | z | p | FDR-p | rg | se | z | p | FDR-p |
| CHARGE IMTmean 2018 | **0.82** | **0.07** | **11.1601** | **6.39E-29** | **1.53E-27** | **0.79** | **0.09** | **8.56** | **1.16E-17** | **2.79E-16** |
| CHARGE IMTmean 2011 | **0.77** | **0.08** | **9.7403** | **2.03E-22** | **2.44E-21** | **0.74** | **0.10** | **7.6706** | **1.71E-14** | **2.05E-13** |
| BMI | **0.13** | **0.03** | **3.8486** | **1.19E-04** | **9.50E-04** | **0.22** | **0.04** | **4.9424** | **7.72E-07** | **6.17E-06** |
| Fasting Glucose | **0.26** | **0.07** | **3.73** | **1.91E-04** | **1.15E-03** | **0.50** | **0.14** | **3.6953** | **2.20E-04** | **1.32E-03** |
| Fasting Insulin | **0.30** | **0.10** | **3.0553** | **2.25E-03** | **1.08E-02** | **-0.17** | **0.05** | **-3.5411** | **3.99E-04** | **1.91E-03** |
| Type 2 Diabetes | **0.21** | **0.08** | **2.8074** | **4.99E-03** | **2.00E-02** | **0.28** | **0.09** | **3.2499** | **1.15E-03** | **4.62E-03** |
| HDL | -0.09 | 0.04 | -2.3216 | 2.03E-02 | 6.94E-02 | **0.17** | **0.06** | **2.823** | **4.76E-03** | **1.63E-02** |
| WHR adjBMI | 0.10 | 0.05 | 2.0435 | 4.10E-02 | 1.23E-01 | **0.24** | **0.09** | **2.7267** | **6.40E-03** | **1.92E-02** |
| Rheumatoid arthritis | 0.11 | 0.06 | 1.7488 | 8.03E-02 | 2.14E-01 | **0.31** | **0.12** | **2.5252** | **1.16E-02** | **3.08E-02** |
| Smoking onset | -0.15 | 0.15 | -0.977 | 3.29E-01 | 5.63E-01 | -0.09 | 0.08 | -1.0276 | 3.04E-01 | 5.21E-01 |
| Fasting Proinsulin | 0.13 | 0.14 | 0.9269 | 3.54E-01 | 5.66E-01 | -0.08 | 0.08 | -0.9862 | 3.24E-01 | 5.18E-01 |
| DBP | 0.02 | 0.02 | 1.1779 | 2.39E-01 | 5.73E-01 | 0.10 | 0.05 | 1.9659 | 4.93E-02 | 1.18E-01 |
| Cholesterol | -0.06 | 0.06 | -0.9904 | 3.22E-01 | 5.94E-01 | -0.07 | 0.06 | -1.1716 | 2.41E-01 | 4.46E-01 |
| Triglycerides | 0.03 | 0.04 | 0.8345 | 4.04E-01 | 6.06E-01 | -0.15 | 0.17 | -0.8927 | 3.72E-01 | 5.58E-01 |
| SBP | 0.02 | 0.02 | 0.7862 | 4.32E-01 | 6.10E-01 | 0.06 | 0.08 | 0.8107 | 4.18E-01 | 5.89E-01 |
| Fasting Insulin adjBMI | 0.10 | 0.10 | 1.001 | 3.17E-01 | 6.34E-01 | 0.10 | 0.08 | 1.3348 | 1.82E-01 | 3.64E-01 |
| Ever Smoker | 0.06 | 0.06 | 1.0029 | 3.16E-01 | 6.89E-01 | 0.18 | 0.13 | 1.3399 | 1.80E-01 | 3.93E-01 |
| Ulcerative Collitis | -0.05 | 0.08 | -0.6214 | 5.34E-01 | 7.12E-01 | 0.02 | 0.02 | 0.7077 | 4.79E-01 | 6.39E-01 |
| Cigarettes per day | 0.05 | 0.11 | 0.465 | 6.42E-01 | 8.11E-01 | 0.10 | 0.17 | 0.5961 | 5.51E-01 | 6.96E-01 |
| LDL | -0.01 | 0.05 | -0.2795 | 7.80E-01 | 8.14E-01 | 0.01 | 0.02 | 0.4633 | 6.43E-01 | 6.71E-01 |
| Irritable Bowel Disorder | -0.02 | 0.05 | -0.3614 | 7.18E-01 | 8.20E-01 | 0.08 | 0.15 | 0.5662 | 5.71E-01 | 6.53E-01 |
| Cardiovascular disease | -0.07 | 0.21 | -0.307 | 7.59E-01 | 8.28E-01 | -0.03 | 0.07 | -0.5171 | 6.05E-01 | 6.60E-01 |
| Crohns Disease | 0.02 | 0.06 | 0.3635 | 7.16E-01 | 8.60E-01 | -0.16 | 0.29 | -0.574 | 5.66E-01 | 6.79E-01 |
| 2hr Glucose | -0.02 | 0.12 | -0.1734 | 8.62E-01 | 8.62E-01 | -0.01 | 0.06 | -0.1364 | 8.92E-01 | 8.92E-01 |

In men, similar to the sex-combined results, IMTmean was positively genetically correlated with the CHARGE IMTmean meta-analyses, total obesity, fasting glucose and fasting insulin. In women, only the genetic correlations with the CHARGE meta-analyses survive FDR correction (STable 5).

### Datamining

The GWAS catalogue was explored (using lead SNPs and locus positions, as per Table 3) to identify previous associations with the cIMT-associated loci (STable 6). Chr7 and Chr8 (6.4Mb) have previously been associated with diastolic blood pressure (DBP), whilst Chr11, Chr13 and both Chr19 loci have previously been associated with coronary artery disease and/or pulse pressure or cIMT. Atherosclerosis is considered a condition requiring both fatty deposits and inflammation in the vascular wall. Therefore it was interesting to note the reported associations of Chr8 (6.4Mb) and Chr11 with immune response or immune components as well as the Chr19 (45Mb) locus with lipid (and other biomarker) levels and Chr7 with fatty liver disease. The Chr19 locus includes the apolipoprotein gene cluster (including *APOE*), which has been the focus of much research into lipid levels, cardiovascular disease and Alzheimer’s disease. In contrast, previous associations at the Chr5, Chr8 (124Mb) and Chr16 loci have no obvious relevance to cIMT. For the vast majority of previous associations with relevant traits, the effects on cIMT were in the expected direction: the alleles associated with increased DBP, systolic blood pressure (SBP), pulse pressure, total cholesterol, low-density lipoprotein (LDL), apolipoprotein E (APOE) and lipoprotein A (LP(a)) levels and lipoprotein phospholipase A2 activity were associated with increased cIMT, as were the alleles associated with decreased HDL levels. The only unexpected finding was that the rs7412 allele associated with decreasing triglyceride (TG) levels was associated with increased cIMT, however the authors suggest that the skewed distribution of lipids in those homozygous for this allele might cause inflation of test statistics ^20^ and that this should be considered when interpreting results.

Of SNPs within significantly associated loci showing at least suggestive evidence of association with IMTmean, 13 SNPs were predicted ^18^ to have functional or coding effects (STable 7). Lead SNPs and those predicted by VEP to have functional or coding effects were assessed for evidence of genotype-specific effects on gene expression levels (eQTLs) in the GTEx dataset (STable 8). Only rs2019090 on Chr11 demonstrated eQTLs in a tissue of obvious relevance, namely the aorta, where the cIMT-increasing allele was associated with increased *RP11-563P16.1* levels. Lack of knowledge about this genes role precludes interpretation of this finding.

### Candidate genes

Of the genes highlighted by eQTL analysis (though in a tissue of unclear relevance), *APOE* (Chr19) has been widely studied in cardiovascular (and other, notably Alzheimer’s) diseases ^21^, whilst the function of *MCPH1-AS1(CTD-2541M15.3)* (Chr8) is thought to be as a regulator of *MCPH1*, a DNA damage response gene with no obvious role in vessel wall biology. Similarly, *CBFA2T3* (Chr16) has documented roles in cancer biology as a transcriptional repressor but again there is no obvious role in vascular biology.

Only *VCAN* (Chr5, women-only locus) is potentially an interesting candidate gene. Its encoded product, versican, is a chondroitin sulphate proteoglycan present in the adventitia and intima of normal blood vessels (reviewed in ^22^). Versican protein levels have been shown to increase dramatically during progression of vascular diseases including atherosclerosis ^22^. Versican exists as a variety of protein isoforms of different sizes, due to alternative splicing, a variety of post-translational modifications and proteolytic cleavage ^22^. The diverse effects of versican in vascular pathology are likely determined by the balance of different sized isoforms and partners in complexes ^22^: larger molecules are better able to bind and thus retain LDL in the vessel wall; loss of the largest versican isoform has been observed in aortic aneurysms; various cytokines promote different sized versican entities or versican degradation; smaller versican isoforms could act directly as mitogens for arterial smooth muscle cells; and the smallest isoform of versican influences elastic fibre formation and therefore vessel function. Therefore, it is interesting to note that the lead SNP for the women-only locus is an eQTL for a versican antisense molecule, *VSCAN1-AS1*, in testes, suggesting yet another mechanism regulating versican expression, although the role of versican in the testes is unknown. Full understanding of versican biology is required before the impact of this SNP is predictable.

## Discussion

Here we present results from a GWAS of the largest single study of IMT to date, and the first sex-specific analysis of IMT. We identified 10 loci (4 of which were novel) associated with cIMT and one locus specific for IMTmean women. We also found genetic correlations with obesity, glucometabolic and lipid traits, which suggest differences between sexes.

Many studies have reported the utility of cIMT measurements as predictors of CVD (reviewed by Katakami et al ^23^), although whether predictive power is independent of traditional risk factors is still unclear. A large part of the discrepancies in the literature is likely due to the different protocols used (which part of the carotid artery is measured, whether plaque is included or not, whether mean (IMTmean) or maximum (IMTmax) values are used) in the analyses. Indeed it has been reported that IMTmean and IMTmax differ in their predictive value ^23^. Therefore the partial overlap in loci associated with IMTmean and IMTmax is of interest. For the most part, SNPs influencing one phenotype also influence the other, but the relative importance of each locus (and/or mechanism) differs. This hints at differences in biology between the 2 measures, which is perhaps not surprising: overall increases in vascular wall thickness could indicate general vascular dysfunction, which is a component of high blood pressure for example. Local increases in vascular wall thickness are likely indicative of plaque formation, which precede vessel occlusion and ischemic events.

Sex-specific differences in cardiovascular disease are well established, therefore the distinctly different IMTmean GWAS results for men compared to women were intriguing. The existence of a women-only locus, whereas the men-only analysis largely resembled the sex-combined analysis, is unexpected. It suggests that the sex-combined results could be “driven” by the men. This is unlikely given the fairly balanced sex distribution of the sample (48% men vs 52% women), however the larger variability in measures in men compared to women could cause this effect. It has been observed that sex-differences in IMT measures can be attributed to sex-differences in risk factors such as BMI and blood pressure ^24^. Despite UKB being approximately 20 years older than the subjects in that study, similar sex differences in risk factors are observed here. It is also worth noting that menopause status is associated both with cIMT measures and cardiovascular risk factors ^25, 26^.

Menopause status was not considered in the study, as direction of effect is unclear. However this factor would be worth considering for future studies of sex differences. Another noteworthy finding is the stronger effect of APOE rs7412 in men compared to women (Beta=0.027 vs 0.016 respectively). To our knowledge this is the first demonstration of a sex effect of APOE variation in a cardiovascular disease phenotype, which is of value when considering the relatively nascent concept of an ageing, sex and *APOE* triad ^27^. Sex-specific differences in APOE-e genotypes associations with Alzheimer’s disease (AD) have been noted (summarised by Fisher et al ^28^), with female carriers of APOE-e3/e4 demonstrating increased risk of AD and faster decline than their male counterparts. Indeed this finding (a variant that increases risk of atherosclerosis for men and AD for women) supports the hypothesis that women have higher rates for AD than men, at least partly because men with CVD die earlier than women, therefore in the at-risk age-range for AD, men have lower CVD risk than women ^28^.

In line with known epidemiology and previous genetic findings ^29^, the genetic correlations between IMTmean or IMTmax and obesity and glucometabolic traits are unsurprising. The lack of genetic correlations with lipid traits is unexpected, but may reflect the relative health, adherence to statin therapy or healthy diet of the general population sample of UKB.

In comparison with the CHARGE consortium IMT meta-analysis, this study was smaller (22,000 vs 68,000) but used a phenotype that was recorded in a consistent manner in all individuals, reducing heterogeneity in measurements. This is likely the reason for identifying loci not reported by CHARGE as well as the larger effect sizes reported here, and is in line with previous observations (for example the C4D ^30^ and CARDIoGRAM findings ^31^).

It should be noted that there is some evidence of inflation of the GWAS statistics, for which there are a number of possible explanations. Polygenicity is an obvious explanation and highly plausible, as it is observed most complex traits. Whilst the pilot phase cIMT data (N∼2500) underwent manual QC procedures, the full IMT dataset (N∼22,000) did not. Comparisons between the QCed and non-QCed data in the pilot study show very high correlation (IMTmean rho=1.0, IMTmax rho=0.98), however the possibility of noise in measurements cannot be excluded. Despite this limitation, the consistent effect sizes and directions of most previous IMT loci suggests that this study is valid.

In conclusion, we have conducted a GWAS of cIMT (mean and maximum measures) in the largest single study sample to date, in which we identified four novel loci and validated 7 of 11 previously reported loci. We also report the first sex-specific analysis of IMT, in which we identified a novel women-only locus. Genetic correlations with obesity and glucometabolic traits were observed, and *VACN* was highlighted as a plausible candidate gene in the women-only locus. Overall, our findings represent an important stimulus for the further elucidation of mechanisms of vascular pathology, particularly differences between men and women and could contribute to stratified medicine approaches.

## Data availability

GWAS summary statistics are available upon request, to the corresponding author. Raw data and coding can be requested via UK Biobank directly.

## Acknowledgements

We thank all participants and staff of the UK Biobank study.

---

**Supplementary Table 1:** Comparison of GWAS significant SNPs for IMTmean on IMTmax

| CHR | BP | SNP | A1 | A0 | A1freq | Beta | IMTmean |  | Beta | IMTmax |  |
| --- | --- | --- | --- | --- | --- | --- | --- | --- | --- | --- | --- |
|  |  |  |  |  |  |  | se | P |  | se | P |
| 5 | 81634499 | rs758080886 | CA | C | 0.15 | 0.013 | 0.002 | **3.10E-08** | 0.015 | 0.003 | 2.10E-07 |
| 5 | 81616381 | rs420800 | G | A | 0.95 | -0.017 | 0.003 | **4.30E-08** | -0.018 | 0.004 | 9.20E-06 |
| 5 | 81618075 | rs224803 | C | T | 0.95 | -0.017 | 0.003 | **4.80E-08** | -0.018 | 0.004 | 1.00E-05 |
| 5 | 81683739 | rs224805 | A | G | 0.05 | 0.017 | 0.003 | **4.10E-08** | 0.017 | 0.004 | 2.50E-05 |
| 5 | 81699253 | rs224960 | A | G | 0.52 | 0.008 | 0.001 | **4.90E-08** | 0.008 | 0.002 | 8.00E-06 |
| 7 | 35423859 | rs2541899 | C | T | 0.24 | 0.009 | 0.002 | **2.20E-08** | 0.009 | 0.002 | 1.70E-05 |
| 7 | 35428513 | rs12155459 | A | T | 0.77 | -0.011 | 0.002 | **2.40E-10** | -0.013 | 0.002 | **2.80E-09** |
| 7 | 35429465 | rs11766731 | G | T | 0.77 | -0.011 | 0.002 | **1.60E-10** | -0.013 | 0.002 | **1.90E-09** |
| 7 | 35435835 | rs2541905 | T | C | 0.23 | 0.011 | 0.002 | **4.60E-10** | 0.013 | 0.002 | **2.50E-09** |
| 7 | 35436932 | rs70981917 | C | CT | 0.43 | 0.008 | 0.001 | **9.70E-09** | 0.009 | 0.002 | **2.70E-06** |
| 7 | 35438779 | rs2013385 | A | G | 0.32 | 0.009 | 0.002 | **5.10E-10** | 0.011 | 0.002 | **2.00E-08** |
| 7 | 35438831 | rs741408 | T | C | 0.23 | 0.011 | 0.002 | **3.80E-11** | 0.013 | 0.002 | **7.50E-10** |
| 7 | 35440055 | rs2592856 | C | T | 0.23 | 0.011 | 0.002 | **2.60E-11** | 0.013 | 0.002 | **5.50E-10** |
| 7 | 35440155 | rs11762074 | T | C | 0.77 | -0.011 | 0.002 | **1.80E-11** | -0.014 | 0.002 | **4.30E-10** |
| 7 | 35446189 | rs2592855 | T | C | 0.23 | 0.011 | 0.002 | **2.40E-11** | 0.013 | 0.002 | **5.20E-10** |
| 7 | 35448018 | rs2250212 | A | G | 0.32 | 0.010 | 0.002 | **3.10E-10** | 0.011 | 0.002 | **1.20E-08** |
| 7 | 35449334 | rs998652 | A | G | 0.23 | 0.011 | 0.002 | **2.60E-11** | 0.013 | 0.002 | **5.70E-10** |
| 7 | 35451830 | rs17765314 | G | A | 0.77 | -0.011 | 0.002 | **2.50E-11** | -0.013 | 0.002 | **5.70E-10** |
| 7 | 35451910 | rs2592851 | T | C | 0.23 | 0.011 | 0.002 | **2.50E-11** | 0.013 | 0.002 | **5.60E-10** |
| 7 | 35452092 | rs1362655 | G | A | 0.23 | 0.011 | 0.002 | **2.60E-11** | 0.013 | 0.002 | **5.90E-10** |
| 7 | 35455088 | rs2592850 | T | A | 0.32 | 0.010 | 0.002 | **2.80E-10** | 0.011 | 0.002 | **1.30E-08** |
| 7 | 35456691 | rs2592849 | A | T | 0.32 | 0.010 | 0.002 | **3.20E-10** | 0.011 | 0.002 | **1.40E-08** |
| 7 | 35457954 | 7:35457954_AT_A | AT | A | 0.32 | 0.009 | 0.002 | **5.50E-10** | 0.012 | 0.002 | **3.90E-09** |
| 7 | 35458137 | rs140476889 | A | AAAAC | 0.32 | 0.010 | 0.002 | **2.90E-10** | 0.011 | 0.002 | **1.20E-08** |
| 7 | 35458551 | rs200832044 | GT | G | 0.32 | 0.009 | 0.002 | **9.00E-10** | 0.011 | 0.002 | **3.20E-08** |
| 7 | 35458934 | rs342976 | T | C | 0.32 | 0.009 | 0.002 | **4.40E-10** | 0.011 | 0.002 | **1.90E-08** |
| 7 | 35459888 | rs342977 | G | A | 0.23 | 0.011 | 0.002 | **4.40E-11** | 0.013 | 0.002 | **1.10E-09** |
| 7 | 35460547 | rs342979 | C | T | 0.32 | 0.009 | 0.002 | **3.90E-10** | 0.011 | 0.002 | **1.60E-08** |
| 7 | 35461538 | rs342981 | G | A | 0.32 | 0.010 | 0.002 | **3.30E-10** | 0.011 | 0.002 | **1.60E-08** |
| 7 | 35461715 | rs2551797 | T | C | 0.32 | 0.010 | 0.002 | **3.30E-10** | 0.011 | 0.002 | **1.60E-08** |
| 7 | 35462145 | rs342982 | G | T | 0.32 | 0.010 | 0.002 | **3.20E-10** | 0.011 | 0.002 | **1.60E-08** |
| 7 | 35464620 | rs34502079 | A | AT | 0.33 | 0.010 | 0.002 | **8.30E-11** | 0.012 | 0.002 | **3.00E-09** |
| 7 | 35464784 | rs342983 | T | G | 0.27 | 0.010 | 0.002 | **1.70E-10** | 0.012 | 0.002 | **3.20E-08** |
| 7 | 35466464 | rs342986 | G | A | 0.23 | 0.011 | 0.002 | **3.90E-11** | 0.013 | 0.002 | **1.00E-09** |
| 7 | 35466658 | rs35361607 | A | AC | 0.37 | 0.008 | 0.001 | **2.80E-08** | 0.009 | 0.002 | 1.80E-06 |
| 7 | 35466755 | rs342987 | T | G | 0.23 | 0.011 | 0.002 | **3.70E-11** | 0.013 | 0.002 | **1.00E-09** |
| 7 | 35467026 | rs342988 | C | T | 0.28 | 0.011 | 0.002 | **8.90E-12** | 0.013 | 0.002 | **8.80E-10** |
| 7 | 35467677 | 7:35467677_TAA_T | TAA | T | 0.24 | 0.011 | 0.002 | **3.80E-11** | 0.013 | 0.002 | **1.00E-09** |
| 7 | 35467896 | rs342989 | A | G | 0.23 | 0.011 | 0.002 | **3.60E-11** | 0.013 | 0.002 | **9.80E-10** |
| 7 | 35468763 | rs17697812 | T | C | 0.77 | -0.011 | 0.002 | **4.90E-11** | -0.013 | 0.002 | **1.20E-09** |
| 7 | 35469045 | rs342990 | G | T | 0.38 | 0.008 | 0.001 | **2.80E-08** | 0.009 | 0.002 | 8.50E-07 |
| 7 | 35469691 | rs342991 | G | T | 0.32 | 0.009 | 0.002 | **8.20E-10** | 0.011 | 0.002 | **2.60E-08** |
| 7 | 35469835 | rs342992 | C | T | 0.23 | 0.011 | 0.002 | **4.90E-11** | 0.013 | 0.002 | **1.20E-09** |
| 7 | 35469898 | rs342993 | G | A | 0.32 | 0.009 | 0.002 | **7.20E-10** | 0.011 | 0.002 | **2.50E-08** |
| 7 | 35471979 | rs342996 | G | C | 0.33 | 0.009 | 0.002 | **6.00E-10** | 0.011 | 0.002 | **2.10E-08** |
| 7 | 35474622 | rs342997 | T | C | 0.32 | 0.009 | 0.002 | **7.20E-10** | 0.011 | 0.002 | **2.60E-08** |
| 7 | 35475980 | rs342998 | T | C | 0.32 | 0.009 | 0.002 | **7.20E-10** | 0.011 | 0.002 | **2.60E-08** |
| 7 | 35477175 | rs343000 | C | T | 0.23 | 0.011 | 0.002 | **5.60E-11** | 0.013 | 0.002 | **1.50E-09** |
| 7 | 35478327 | rs343001 | C | G | 0.32 | 0.009 | 0.002 | **7.30E-10** | 0.011 | 0.002 | **2.60E-08** |
| 7 | 35478515 | rs343002 | T | C | 0.23 | 0.011 | 0.002 | **5.60E-11** | 0.013 | 0.002 | **1.50E-09** |
| 7 | 35479439 | rs343003 | C | T | 0.23 | 0.011 | 0.002 | **5.90E-11** | 0.013 | 0.002 | **1.50E-09** |
| 7 | 35481642 | rs343004 | C | A | 0.32 | 0.009 | 0.002 | **9.00E-10** | 0.011 | 0.002 | **3.00E-08** |
| 7 | 35482940 | rs343005 | G | T | 0.32 | 0.009 | 0.002 | **8.70E-10** | 0.011 | 0.002 | **2.60E-08** |
| 7 | 35483018 | rs343006 | C | A | 0.32 | 0.009 | 0.002 | **9.10E-10** | 0.011 | 0.002 | **2.60E-08** |
| 7 | 35483055 | rs343007 | G | T | 0.32 | 0.009 | 0.002 | **8.00E-10** | 0.011 | 0.002 | **2.60E-08** |
| 7 | 35485648 | rs343010 | C | A | 0.32 | 0.009 | 0.002 | **7.50E-10** | 0.011 | 0.002 | **2.60E-08** |
| 7 | 35487040 | 7:35487040_CT_C | CT | C | 0.29 | 0.011 | 0.002 | **3.20E-09** | 0.013 | 0.002 | **1.60E-08** |
| 7 | 35488349 | rs343011 | T | G | 0.23 | 0.011 | 0.002 | **5.50E-11** | 0.013 | 0.002 | **1.50E-09** |
| 7 | 35488769 | rs343012 | C | A | 0.32 | 0.009 | 0.002 | **7.40E-10** | 0.011 | 0.002 | **2.70E-08** |
| 7 | 35491532 | rs343016 | G | C | 0.32 | 0.009 | 0.002 | **8.00E-10** | 0.011 | 0.002 | **2.60E-08** |
| 7 | 35492168 | rs343019 | G | C | 0.41 | 0.009 | 0.001 | **7.10E-10** | 0.010 | 0.002 | 1.60E-07 |
| 7 | 35493178 | rs343020 | T | C | 0.32 | 0.009 | 0.002 | **8.20E-10** | 0.011 | 0.002 | **2.40E-08** |
| 7 | 35493309 | rs343021 | T | C | 0.32 | 0.009 | 0.002 | **8.10E-10** | 0.011 | 0.002 | **2.40E-08** |
| 7 | 35493603 | rs343022 | C | T | 0.32 | 0.009 | 0.002 | **8.70E-10** | 0.011 | 0.002 | **2.20E-08** |
| 7 | 35494013 | rs343024 | C | G | 0.41 | 0.009 | 0.001 | **5.80E-10** | 0.010 | 0.002 | 1.20E-07 |
| 7 | 35494243 | rs777980 | G | A | 0.32 | 0.009 | 0.002 | **6.10E-10** | 0.011 | 0.002 | **1.70E-08** |
| 7 | 35494556 | rs702843 | G | C | 0.41 | 0.009 | 0.001 | **3.90E-10** | 0.010 | 0.002 | **7.30E-08** |
| 7 | 35495599 | rs343025 | T | A | 0.37 | 0.009 | 0.001 | **5.00E-09** | 0.010 | 0.002 | 2.80E-07 |
| 7 | 35496678 | rs343026 | T | G | 0.32 | 0.009 | 0.002 | **5.20E-10** | 0.011 | 0.002 | **1.50E-08** |
| 7 | 35497055 | rs35099106 | C | CT | 0.23 | 0.011 | 0.002 | **3.70E-11** | 0.013 | 0.002 | **8.70E-10** |
| 7 | 35498200 | rs343029 | A | G | 0.23 | 0.011 | 0.002 | **3.50E-11** | 0.013 | 0.002 | **8.90E-10** |
| 7 | 35500493 | rs343031 | G | A | 0.32 | 0.009 | 0.002 | **5.00E-10** | 0.011 | 0.002 | **1.50E-08** |
| 7 | 35501883 | rs343034 | C | T | 0.37 | 0.009 | 0.001 | **5.20E-09** | 0.010 | 0.002 | 2.70E-07 |
| 7 | 35503445 | rs343036 | G | C | 0.23 | 0.011 | 0.002 | **3.50E-11** | 0.013 | 0.002 | **8.70E-10** |
| 7 | 35503600 | rs343037 | G | T | 0.23 | 0.011 | 0.002 | **3.60E-11** | 0.013 | 0.002 | **8.70E-10** |
| 7 | 35503696 | rs343038 | G | C | 0.32 | 0.009 | 0.002 | **4.70E-10** | 0.011 | 0.002 | **1.40E-08** |
| 7 | 35505204 | rs343039 | C | T | 0.23 | 0.011 | 0.002 | **3.90E-11** | 0.013 | 0.002 | **7.90E-10** |
| 7 | 35508859 | rs343044 | A | G | 0.20 | 0.010 | 0.002 | **3.60E-08** | 0.012 | 0.002 | 4.30E-07 |
| 7 | 35509024 | rs343045 | C | A | 0.20 | 0.010 | 0.002 | **3.60E-08** | 0.012 | 0.002 | 4.40E-07 |
| 7 | 35515295 | rs343048 | G | A | 0.20 | 0.010 | 0.002 | **3.30E-08** | 0.012 | 0.002 | 4.20E-07 |
| 7 | 35519519 | rs343052 | G | T | 0.20 | 0.010 | 0.002 | **3.50E-08** | 0.012 | 0.002 | 4.20E-07 |
| 7 | 35522553 | rs343053 | A | G | 0.17 | 0.011 | 0.002 | **4.30E-09** | 0.012 | 0.002 | 4.60E-07 |
| 7 | 35523883 | rs343055 | G | A | 0.17 | 0.011 | 0.002 | **3.50E-09** | 0.013 | 0.002 | 2.90E-07 |
| 7 | 35526022 | rs1646989 | T | G | 0.20 | 0.010 | 0.002 | **1.50E-08** | 0.012 | 0.002 | 1.60E-07 |
| 7 | 35526024 | rs1646988 | T | A | 0.20 | 0.010 | 0.002 | **1.50E-08** | 0.012 | 0.002 | 1.60E-07 |
| 7 | 35526390 | rs343056 | A | G | 0.20 | 0.010 | 0.002 | **7.00E-09** | 0.012 | 0.002 | 1.30E-07 |
| 7 | 35528567 | rs11766156 | G | C | 0.80 | -0.010 | 0.002 | **7.40E-09** | -0.012 | 0.002 | 1.20E-07 |
| 7 | 35532991 | rs73087411 | C | T | 0.82 | -0.010 | 0.002 | **4.60E-08** | -0.011 | 0.002 | 2.80E-06 |
| 8 | 6470650 | rs3020257 | C | A | 0.76 | 0.010 | 0.002 | **6.60E-10** | 0.009 | 0.002 | 8.10E-05 |
| 8 | 6471126 | rs2980664 | G | A | 0.76 | 0.011 | 0.002 | **2.50E-10** | 0.009 | 0.002 | 2.60E-05 |
| 8 | 6471182 | rs3020258 | C | T | 0.78 | 0.010 | 0.002 | **2.00E-09** | 0.009 | 0.002 | 9.60E-05 |
| 8 | 6471995 | rs2959802 | G | A | 0.42 | -0.008 | 0.001 | **3.30E-08** | -0.007 | 0.002 | 9.20E-05 |
| 8 | 6472105 | rs2959801 | G | A | 0.69 | 0.010 | 0.002 | **2.00E-10** | 0.009 | 0.002 | 9.10E-06 |
| 8 | 6472603 | rs11137038 | G | A | 0.64 | 0.010 | 0.001 | **6.30E-11** | 0.008 | 0.002 | 2.00E-05 |
| 8 | 6472721 | rs12543564 | T | C | 0.67 | 0.009 | 0.002 | **1.30E-08** | 0.009 | 0.002 | 1.70E-05 |
| 8 | 6472795 | rs3020259 | C | A | 0.65 | 0.009 | 0.001 | **2.30E-10** | 0.008 | 0.002 | 2.10E-05 |
| 8 | 6473118 | rs11998083 | C | T | 0.64 | 0.010 | 0.001 | **4.50E-11** | 0.008 | 0.002 | 1.30E-05 |
| 8 | 6473356 | rs1256 | C | G | 0.70 | 0.010 | 0.002 | **1.80E-11** | 0.010 | 0.002 | 1.20E-06 |
| 8 | 6473715 | rs2959799 | A | G | 0.63 | 0.010 | 0.001 | **1.10E-10** | 0.008 | 0.002 | 1.80E-05 |
| 8 | 6473746 | rs2959798 | A | G | 0.63 | 0.009 | 0.001 | **1.20E-10** | 0.008 | 0.002 | 1.80E-05 |
| 8 | 6473758 | rs2454517 | A | G | 0.70 | 0.010 | 0.002 | **4.30E-11** | 0.010 | 0.002 | 2.40E-06 |
| 8 | 6474136 | rs2936501 | A | G | 0.63 | 0.009 | 0.001 | **1.60E-10** | 0.008 | 0.002 | 1.70E-05 |
| 8 | 6475113 | rs2515557 | G | A | 0.70 | 0.009 | 0.002 | **3.90E-08** | 0.008 | 0.002 | 6.60E-05 |
| 8 | 6475489 | rs2980662 | G | C | 0.69 | 0.009 | 0.002 | **2.60E-09** | 0.009 | 0.002 | 1.90E-05 |
| 8 | 6476626 | rs7005662 | C | T | 0.72 | 0.010 | 0.002 | **1.50E-10** | 0.010 | 0.002 | 2.50E-06 |
| 8 | 6478481 | rs2253560 | G | A | 0.72 | 0.010 | 0.002 | **7.80E-11** | 0.009 | 0.002 | 4.20E-06 |
| 8 | 6478960 | rs11137040 | C | G | 0.71 | 0.010 | 0.002 | **7.90E-10** | 0.009 | 0.002 | 2.70E-05 |
| 8 | 6479178 | rs2912016 | C | A | 0.65 | 0.009 | 0.001 | **6.70E-10** | 0.008 | 0.002 | 4.70E-05 |
| 8 | 6479282 | rs11137041 | G | T | 0.65 | 0.009 | 0.001 | **2.20E-09** | 0.008 | 0.002 | 7.30E-05 |
| 8 | 6479432 | rs2515559 | A | G | 0.72 | 0.010 | 0.002 | **5.10E-10** | 0.009 | 0.002 | 8.40E-06 |
| 8 | 6479497 | rs2011423 | G | A | 0.67 | 0.009 | 0.002 | **1.40E-09** | 0.008 | 0.002 | 4.10E-05 |
| 8 | 6483059 | rs201338894 | ATAAAT | A | 0.73 | 0.011 | 0.002 | **9.30E-12** | 0.010 | 0.002 | 1.70E-06 |
| 8 | 6483482 | rs2912057 | C | T | 0.60 | 0.008 | 0.001 | **2.60E-08** | 0.008 | 0.002 | 4.10E-05 |
| 8 | 6484505 | 8:6484505_CA_C | CA | C | 0.72 | 0.011 | 0.002 | **3.10E-12** | 0.010 | 0.002 | 9.40E-07 |
| 8 | 6484510 | rs2454524 | A | C | 0.72 | 0.011 | 0.002 | **3.10E-12** | 0.010 | 0.002 | 9.50E-07 |
| 8 | 6484530 | rs2246801 | C | T | 0.72 | 0.011 | 0.002 | **4.50E-12** | 0.010 | 0.002 | 1.10E-06 |
| 8 | 6484635 | rs2278147 | C | A | 0.71 | 0.011 | 0.002 | **4.10E-13** | 0.010 | 0.002 | 6.10E-07 |
| 8 | 6484982 | rs2448611 | C | T | 0.70 | 0.010 | 0.002 | **1.50E-10** | 0.010 | 0.002 | 1.30E-06 |
| 8 | 6485295 | rs2912062 | C | G | 0.70 | 0.011 | 0.002 | **2.70E-13** | 0.010 | 0.002 | 2.70E-07 |
| 8 | 6486033 | rs2912063 | A | G | 0.71 | 0.011 | 0.002 | **4.20E-13** | 0.010 | 0.002 | 4.80E-07 |
| 8 | 6487449 | rs3020263 | G | A | 0.72 | 0.011 | 0.002 | **4.40E-12** | 0.010 | 0.002 | 1.00E-06 |
| 8 | 6487727 | rs2936511 | C | G | 0.63 | 0.008 | 0.001 | **2.20E-08** | 0.007 | 0.002 | 5.30E-04 |
| 8 | 6488317 | rs569123053 | A | C | 0.80 | 0.011 | 0.002 | **2.70E-09** | 0.010 | 0.002 | 6.50E-05 |
| 8 | 6488505 | rs2911968 | T | C | 0.62 | 0.008 | 0.001 | **2.30E-08** | 0.007 | 0.002 | 4.20E-04 |
| 8 | 6488577 | rs3020264 | G | A | 0.72 | 0.011 | 0.002 | **5.80E-12** | 0.010 | 0.002 | 1.40E-06 |
| 8 | 6488710 | rs2912064 | C | T | 0.65 | 0.009 | 0.001 | **2.60E-10** | 0.008 | 0.002 | 1.10E-04 |
| 8 | 6488787 | rs1974946 | G | A | 0.72 | 0.011 | 0.002 | **5.70E-12** | 0.010 | 0.002 | 1.60E-06 |
| 8 | 6488921 | rs3020265 | G | A | 0.63 | 0.008 | 0.001 | **1.60E-08** | 0.007 | 0.002 | 3.30E-04 |
| 8 | 6488990 | 8:6488990_CA_C | CA | C | 0.62 | 0.008 | 0.001 | **2.40E-08** | 0.007 | 0.002 | 3.60E-04 |
| 8 | 6490200 | rs2433143 | C | T | 0.73 | 0.010 | 0.002 | **6.50E-11** | 0.009 | 0.002 | 3.40E-05 |
| 8 | 6490544 | 8:6490544_TA_T | TA | T | 0.70 | 0.010 | 0.002 | **2.00E-11** | 0.008 | 0.002 | 3.90E-05 |
| 8 | 6490975 | rs2515561 | G | A | 0.66 | 0.009 | 0.001 | **6.70E-10** | 0.007 | 0.002 | 2.60E-04 |
| 8 | 6491181 | rs1834451 | T | C | 0.61 | 0.009 | 0.001 | **1.60E-09** | 0.007 | 0.002 | 2.70E-04 |
| 8 | 6492014 | 8:6492014_CTG_C | CTG | C | 0.67 | 0.008 | 0.002 | **1.90E-08** | 0.006 | 0.002 | 9.60E-04 |
| 8 | 6492391 | rs2980655 | A | T | 0.67 | 0.009 | 0.002 | **6.70E-09** | 0.007 | 0.002 | 4.70E-04 |
| 8 | 6492498 | rs3020266 | G | A | 0.67 | 0.009 | 0.002 | **8.20E-09** | 0.007 | 0.002 | 5.20E-04 |
| 8 | 6492792 | 8:6492792_AC_A | AC | A | 0.66 | 0.009 | 0.002 | **1.10E-08** | 0.007 | 0.002 | 5.00E-04 |
| 8 | 6493910 | rs2042670 | A | C | 0.66 | 0.008 | 0.001 | **1.40E-08** | 0.006 | 0.002 | 1.80E-03 |
| 8 | 6494850 | rs2515563 | G | C | 0.69 | 0.010 | 0.002 | **2.50E-10** | 0.008 | 0.002 | 9.50E-05 |
| 8 | 124579985 | rs6470156 | G | A | 0.73 | -0.011 | 0.002 | **5.30E-13** | -0.009 | 0.002 | 2.10E-05 |
| 8 | 124594184 | rs13270370 | G | A | 0.55 | -0.008 | 0.001 | **2.40E-08** | -0.006 | 0.002 | 2.60E-03 |
| 8 | 124594923 | rs11776999 | G | A | 0.80 | -0.010 | 0.002 | **3.50E-08** | -0.008 | 0.002 | 5.40E-04 |
| 8 | 124599786 | rs13251106 | C | T | 0.80 | -0.010 | 0.002 | **3.60E-08** | -0.008 | 0.002 | 5.60E-04 |
| 8 | 124601535 | rs12547541 | G | A | 0.80 | -0.010 | 0.002 | **3.70E-08** | -0.008 | 0.002 | 5.50E-04 |
| 8 | 124602054 | rs35534969 | A | C | 0.55 | -0.008 | 0.001 | **2.20E-08** | -0.006 | 0.002 | 2.40E-03 |
| 8 | 124607159 | rs34557926 | C | T | 0.64 | -0.011 | 0.001 | **1.40E-13** | -0.008 | 0.002 | 2.60E-05 |
| 8 | 124608162 | rs2891781 | C | T | 0.64 | -0.011 | 0.001 | **2.70E-13** | -0.008 | 0.002 | 3.90E-05 |
| 8 | 124608614 | rs7006122 | T | G | 0.64 | -0.011 | 0.001 | **3.10E-13** | -0.008 | 0.002 | 4.50E-05 |
| 10 | 30138169 | rs7912501 | A | G | 0.30 | 0.009 | 0.002 | **2.40E-08** | 0.007 | 0.002 | 1.20E-03 |
| 10 | 30138411 | rs6481643 | T | C | 0.30 | 0.009 | 0.002 | **2.20E-08** | 0.007 | 0.002 | 1.20E-03 |
| 10 | 30144471 | rs10763762 | T | C | 0.30 | 0.008 | 0.002 | **4.80E-08** | 0.006 | 0.002 | 1.60E-03 |
| 10 | 30161168 | rs1929257 | A | G | 0.31 | 0.009 | 0.002 | **6.00E-09** | 0.007 | 0.002 | 5.70E-04 |
| 10 | 30162423 | rs2150562 | A | G | 0.31 | 0.009 | 0.002 | **8.70E-09** | 0.007 | 0.002 | 5.60E-04 |
| 10 | 30163133 | rs777393148 | TC | T | 0.31 | 0.008 | 0.002 | **3.30E-08** | 0.007 | 0.002 | 1.00E-03 |
| 10 | 30165983 | rs7096778 | T | C | 0.41 | 0.008 | 0.001 | **1.60E-08** | 0.007 | 0.002 | 1.40E-04 |
| 10 | 30166035 | rs7078837 | C | A | 0.41 | 0.008 | 0.001 | **1.40E-08** | 0.007 | 0.002 | 1.10E-04 |
| 10 | 30166547 | rs10826719 | G | A | 0.41 | 0.008 | 0.001 | **1.70E-08** | 0.007 | 0.002 | 1.30E-04 |
| 10 | 30167009 | rs1571759 | T | C | 0.41 | 0.008 | 0.001 | **1.30E-08** | 0.007 | 0.002 | 1.20E-04 |
| 10 | 30167754 | rs10740811 | G | A | 0.41 | 0.008 | 0.001 | **1.10E-08** | 0.007 | 0.002 | 8.50E-05 |
| 10 | 30168031 | rs2369339 | T | C | 0.32 | 0.009 | 0.002 | **5.10E-09** | 0.007 | 0.002 | 2.60E-04 |
| 10 | 30168699 | rs2202 | C | T | 0.32 | 0.009 | 0.002 | **8.80E-09** | 0.007 | 0.002 | 2.80E-04 |
| 10 | 30169653 | rs10763764 | A | T | 0.42 | 0.008 | 0.001 | **1.10E-08** | 0.007 | 0.002 | 8.60E-05 |
| 10 | 30170487 | rs914279 | T | G | 0.42 | 0.008 | 0.001 | **6.80E-09** | 0.007 | 0.002 | 7.00E-05 |
| 11 | 103660567 | rs974819 | T | C | 0.29 | 0.010 | 0.002 | **5.60E-10** | 0.010 | 0.002 | 3.70E-07 |
| 11 | 103668962 | rs2019090 | A | T | 0.29 | 0.010 | 0.002 | **3.70E-10** | 0.010 | 0.002 | 3.30E-07 |
| 11 | 103669291 | rs796784254 | T | TTATTGAA | 0.29 | 0.010 | 0.002 | **5.80E-10** | 0.010 | 0.002 | 4.20E-07 |
| 11 | 103673277 | rs2128739 | A | C | 0.28 | 0.009 | 0.002 | **2.20E-09** | 0.010 | 0.002 | 1.10E-06 |
| 11 | 103673294 | rs2839812 | T | A | 0.28 | 0.009 | 0.002 | **2.20E-09** | 0.010 | 0.002 | 1.10E-06 |
| 11 | 103693627 | rs11226029 | G | A | 0.29 | 0.009 | 0.002 | **1.10E-08** | 0.009 | 0.002 | 3.30E-06 |
| 11 | 103696851 | rs1384705 | C | T | 0.29 | 0.009 | 0.002 | **1.30E-08** | 0.009 | 0.002 | 3.20E-06 |
| 13 | 111049623 | rs9515203 | T | C | 0.74 | -0.009 | 0.002 | **1.10E-08** | -0.009 | 0.002 | 1.20E-05 |
| 16 | 88966667 | rs844396 | C | T | 0.68 | 0.008 | 0.002 | **4.50E-08** | 0.009 | 0.002 | 1.40E-05 |
| 16 | 88968540 | rs564669 | T | C | 0.62 | 0.008 | 0.001 | **4.30E-08** | 0.008 | 0.002 | 2.70E-05 |
| 16 | 88969969 | rs865102 | A | G | 0.65 | 0.008 | 0.001 | **9.10E-09** | 0.008 | 0.002 | 1.30E-05 |
| 16 | 88970040 | rs4995274 | T | C | 0.65 | 0.008 | 0.001 | **3.20E-08** | 0.008 | 0.002 | 2.90E-05 |
| 16 | 88970776 | rs475796 | C | G | 0.65 | 0.008 | 0.001 | **1.00E-08** | 0.008 | 0.002 | 1.40E-05 |
| 16 | 88970818 | rs561795 | A | G | 0.65 | 0.008 | 0.001 | **1.30E-08** | 0.008 | 0.002 | 1.30E-05 |
| 16 | 88972554 | rs488251 | G | A | 0.68 | 0.008 | 0.002 | **3.30E-08** | 0.008 | 0.002 | 4.00E-05 |
| 16 | 88974516 | rs548591 | A | G | 0.62 | 0.008 | 0.001 | **1.80E-08** | 0.008 | 0.002 | 1.60E-05 |
| 16 | 88974860 | rs533406 | A | G | 0.62 | 0.008 | 0.001 | **2.70E-08** | 0.008 | 0.002 | 2.40E-05 |
| 16 | 88975447 | rs8058234 | C | G | 0.62 | 0.008 | 0.001 | **3.20E-08** | 0.008 | 0.002 | 4.10E-05 |
| 16 | 88975910 | rs7405380 | G | C | 0.62 | 0.008 | 0.001 | **2.20E-08** | 0.008 | 0.002 | 3.80E-05 |
| 16 | 88975954 | rs7404039 | A | G | 0.62 | 0.008 | 0.001 | **2.70E-08** | 0.008 | 0.002 | 5.20E-05 |
| 16 | 88976477 | rs1673931 | T | C | 0.62 | 0.008 | 0.001 | **2.30E-08** | 0.008 | 0.002 | 3.50E-05 |
| 16 | 88976968 | rs529900 | G | C | 0.62 | 0.008 | 0.001 | **2.50E-08** | 0.008 | 0.002 | 3.90E-05 |
| 16 | 88982789 | rs12444726 | G | T | 0.63 | 0.008 | 0.001 | **4.70E-08** | 0.007 | 0.002 | 2.90E-04 |
| 16 | 88987861 | rs1673932 | T | C | 0.63 | 0.008 | 0.001 | **3.50E-08** | 0.008 | 0.002 | 5.30E-05 |
| 16 | 88988989 | rs561732 | T | C | 0.65 | 0.008 | 0.001 | **4.70E-08** | 0.008 | 0.002 | 3.80E-05 |
| 19 | 41188985 | rs113228202 | G | A | 0.99 | -0.036 | 0.007 | **3.40E-08** | -0.034 | 0.008 | 6.70E-05 |
| 19 | 41333152 | rs111689747 | G | A | 0.99 | -0.040 | 0.006 | **7.10E-10** | -0.044 | 0.008 | 2.20E-07 |
| 19 | 41371458 | rs111240159 | C | T | 0.99 | -0.037 | 0.006 | **4.00E-09** | -0.041 | 0.008 | 3.60E-07 |
| 19 | 45337737 | rs11667253 | C | T | 0.30 | -0.009 | 0.002 | **2.70E-08** | -0.009 | 0.002 | 1.10E-05 |
| 19 | 45338220 | rs4369782 | A | G | 0.30 | -0.009 | 0.002 | **2.90E-08** | -0.009 | 0.002 | 9.60E-06 |
| 19 | 45338493 | rs4605275 | T | C | 0.30 | -0.009 | 0.002 | **3.40E-08** | -0.009 | 0.002 | 9.20E-06 |
| 19 | 45341948 | rs10407439 | A | G | 0.30 | -0.009 | 0.002 | **2.30E-08** | -0.009 | 0.002 | 9.90E-06 |
| 19 | 45342241 | rs4239533 | A | G | 0.30 | -0.009 | 0.002 | **1.80E-08** | -0.009 | 0.002 | 8.50E-06 |
| 19 | 45397229 | rs1160983 | G | A | 0.97 | 0.023 | 0.004 | **4.70E-08** | 0.030 | 0.005 | **1.70E-08** |
| 19 | 45412079 | rs7412 | C | T | 0.92 | 0.021 | 0.003 | **2.10E-15** | 0.025 | 0.003 | **1.00E-13** |
| 19 | 45413233 | rs1065853 | G | T | 0.92 | 0.021 | 0.003 | **1.60E-15** | 0.026 | 0.003 | **6.90E-14** |
| 19 | 45414399 | rs72654473 | C | A | 0.90 | 0.013 | 0.002 | **2.40E-08** | 0.017 | 0.003 | **8.70E-09** |
| 19 | 45415640 | rs445925 | G | A | 0.89 | 0.013 | 0.002 | **3.20E-08** | 0.017 | 0.003 | **1.30E-08** |
| 19 | 45416831 | rs390082 | T | G | 0.89 | 0.013 | 0.002 | **4.10E-08** | 0.017 | 0.003 | **2.00E-08** |
| 19 | 45425178 | rs190712692 | G | A | 0.95 | 0.023 | 0.003 | **1.30E-12** | 0.028 | 0.004 | **7.80E-11** |
| 19 | 45426792 | rs141622900 | G | A | 0.95 | 0.023 | 0.003 | **1.90E-12** | 0.026 | 0.004 | **4.00E-10** |

**Supplementary Table 2:** Effects of previsouly reported IMT loci in the UKB GWAS

| SNP | Genomic region | Nearest Coding Gene | Alleles Effect/ Other | Effect allele frequency | Franceschini |  | Alleles Effect/ Other | UKB |  |  |  |
| --- | --- | --- | --- | --- | --- | --- | --- | --- | --- | --- | --- |
|  |  |  |  |  | Beta (SE) | p-value |  | A1 freq | beta | P | Consistent |
| rs201648240 | chr1:208953176 | | D/I | 0.83 | -0.0062 (0.0011) | $3.6 \times 10^{-9}$ | | | | | |
| rs224904 | chr5:81637916 | *ATP6AP1L* | C/G | 0.95 | -0.0088 (0.0016) | $5.1 \times 10^{-8}$ | G/C | 0.05 | 0.0163 | 7.20E-07 | y |
| rs6907215 | chr6:143608968 | *AIG1* | T/C | 0.6 | -0.0040 (0.0007) | $4.7 \times 10^{-8}$ | C/T | 0.40 | 0.0060 | 6.80E-05 | Y |
| rs13225723 | chr7:106416467 | *PIK3CG* | A/G | 0.22 | 0.0052 (0.0009) | $3.2 \times 10^{-9}$ | G/A | 0.89 | -0.0097 | 0.0004 | y |
| rs2912063 | chr8:6486033 | *MCPH1* | A/G | 0.71 | 0.0045 (0.0008) | $8.9 \times 10^{-9}$ | A/G | 0.71 | 0.0122 | **8.3E-14** | y |
| rs11785239 | chr8:8205010 | *SGK223* | T/C | 0.65 | -0.0043 (0.0008) | $8.7 \times 10^{-9}$ | C/T | 0.37 | -0.0045 | 0.0033 | n |
| rs11196033 | chr10:114410998 | *VTI1A* | A/C | 0.48 | 0.0042 (0.0008) | $3.9 \times 10^{-8}$ | C/A | 0.49 | 0.0011 | 0.4500 | n |
| rs844396 | chr16:88966667 | *CBFA2T3* | T/C | 0.3 | -0.0051 (0.0009) | $6.0 \times 10^{-9}$ | C/T | 0.68 | 0.0086 | **4.80E-08** | y |
| rs148147734<sup>a</sup> | 8:123401537-indel | *ZHX2* | -/G | 0.54 | 0.0050 (0.0007) | $3 \times 10^{-11}$ | G/A | 0.54 | 0.0015 | 3.70E-05 | y |
| rs200482500 | 8:10606223-indel | *PINX1* | -/GTACC | 0.52 | 0.0056 (0.0008) | $7 \times 10^{-12}$ | | | | | |
| rs7412 | 19:45412079 | *APOE* | T/C | 0.08 | -0.0119 (0.0015) | $1 \times 10^{-14}$ | C/T | | 0.0214 | **7.1E-15** | y |

<sup>a</sup>Published cIMT SNP in LD with our most significant SNP: rs11781551 ( $r^2 = 0.95$ with rs148147734)

**Supplemental Table 3:**
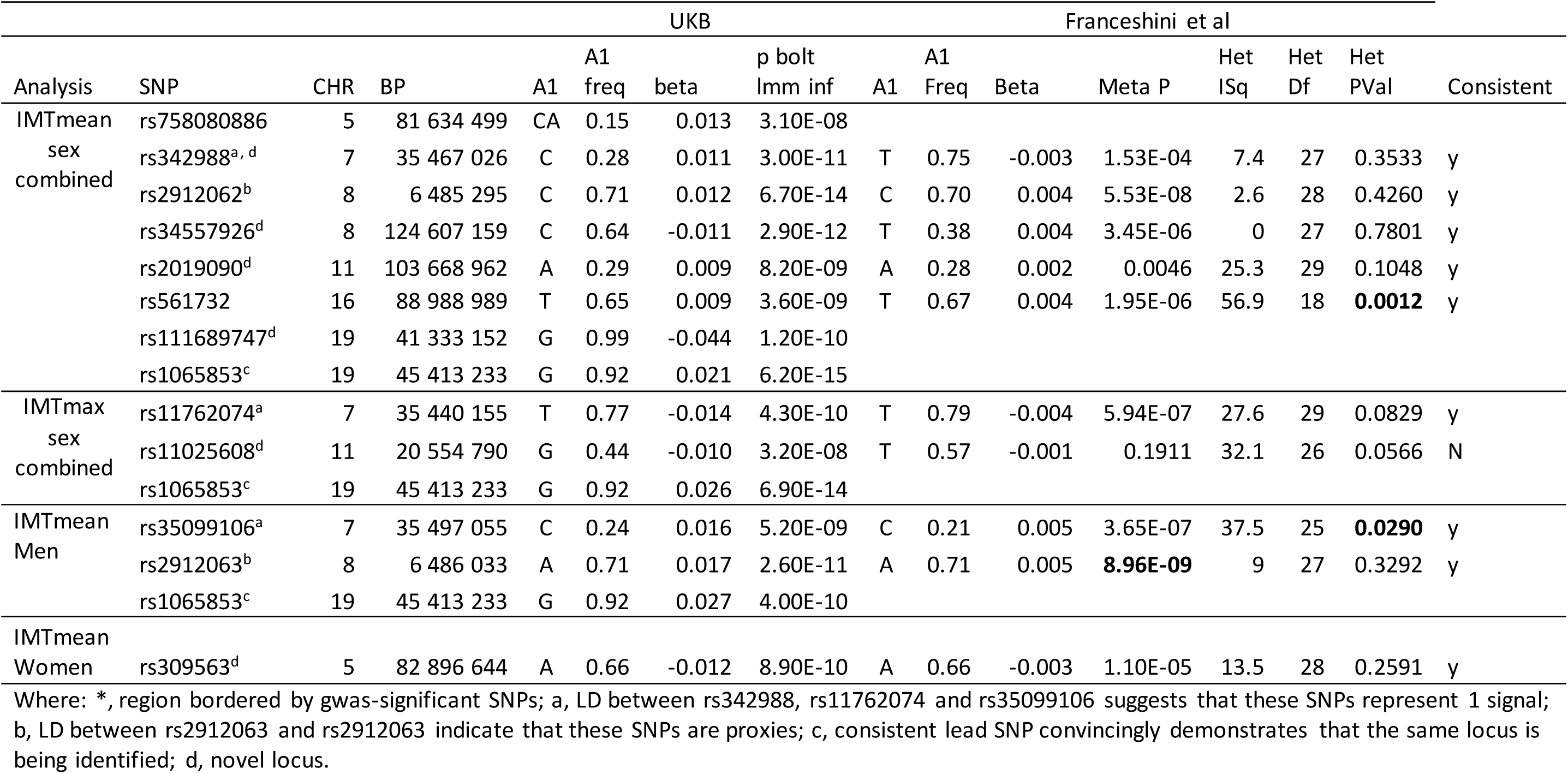
Effects of GWAS-significant loci reported here in the CHARGE meta-analysis of IMT.

**Supplementary Table 4:**
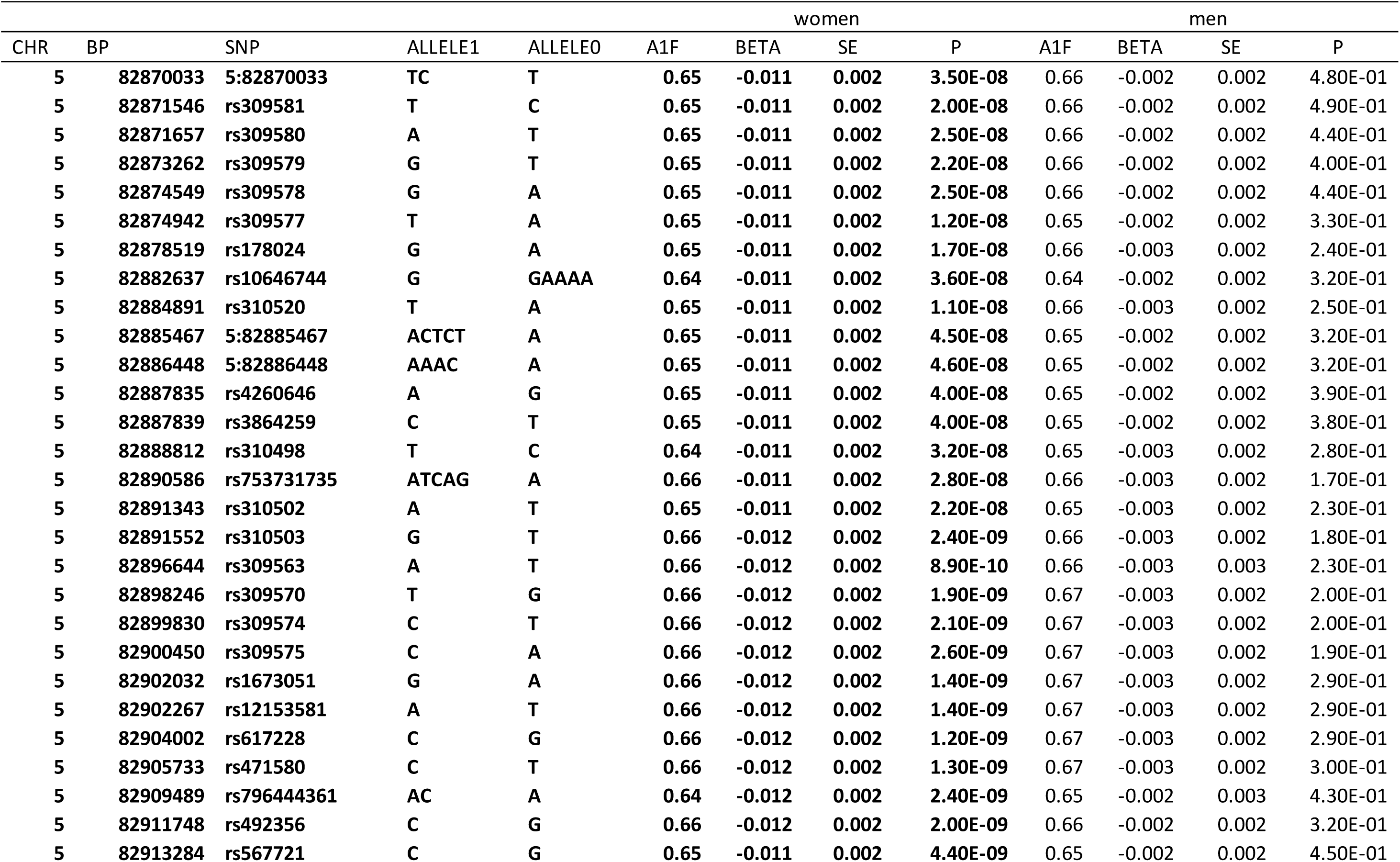
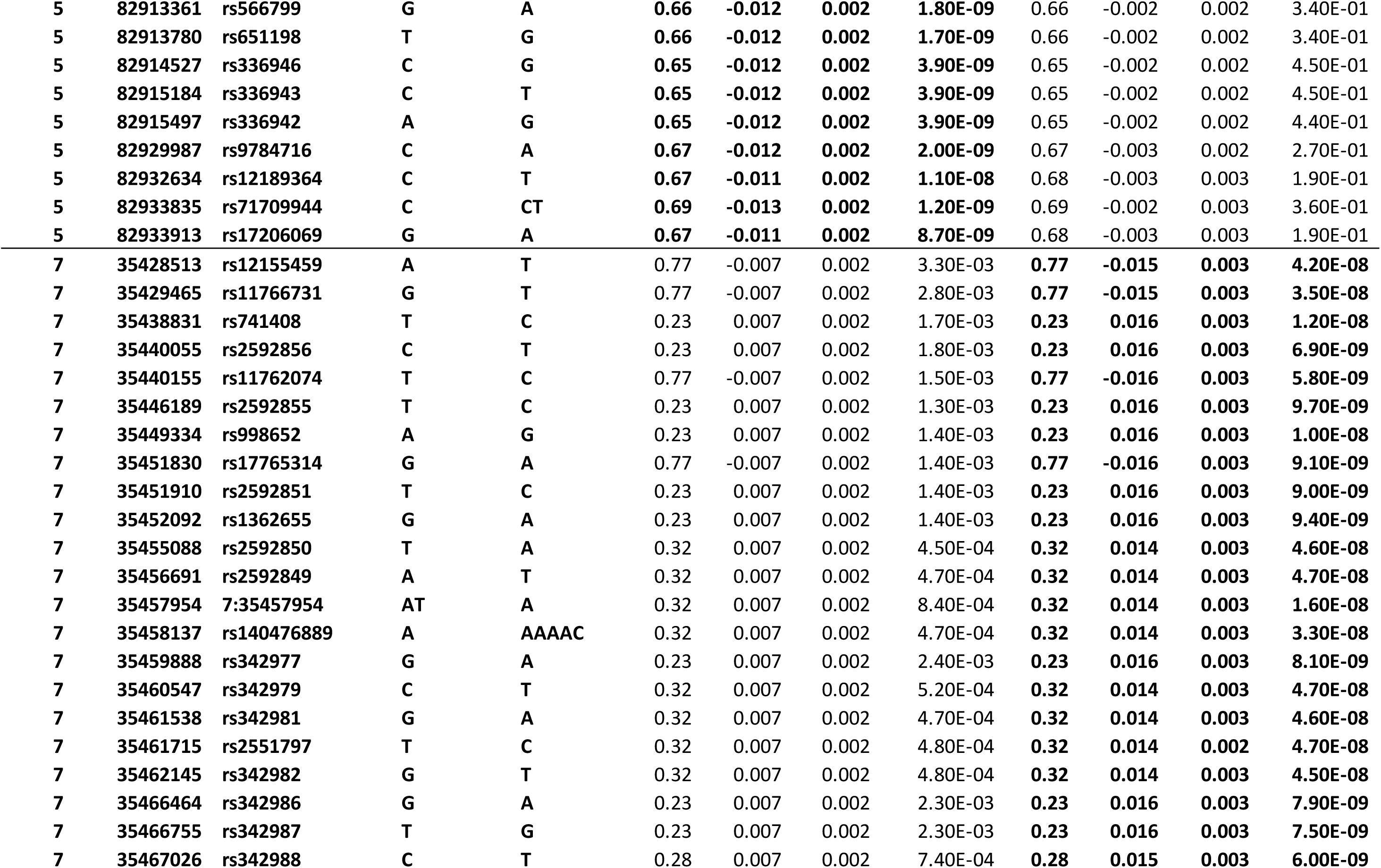
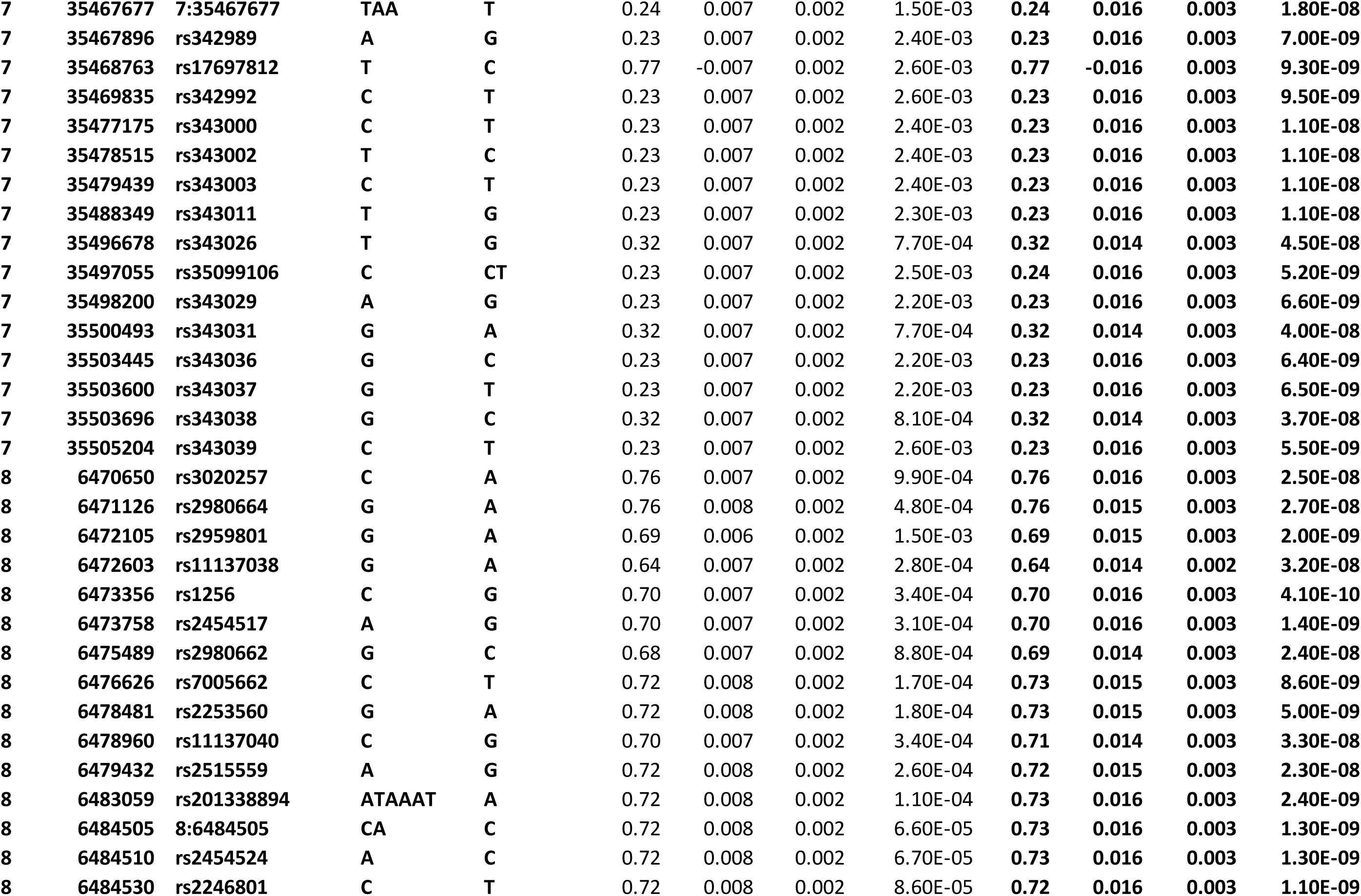
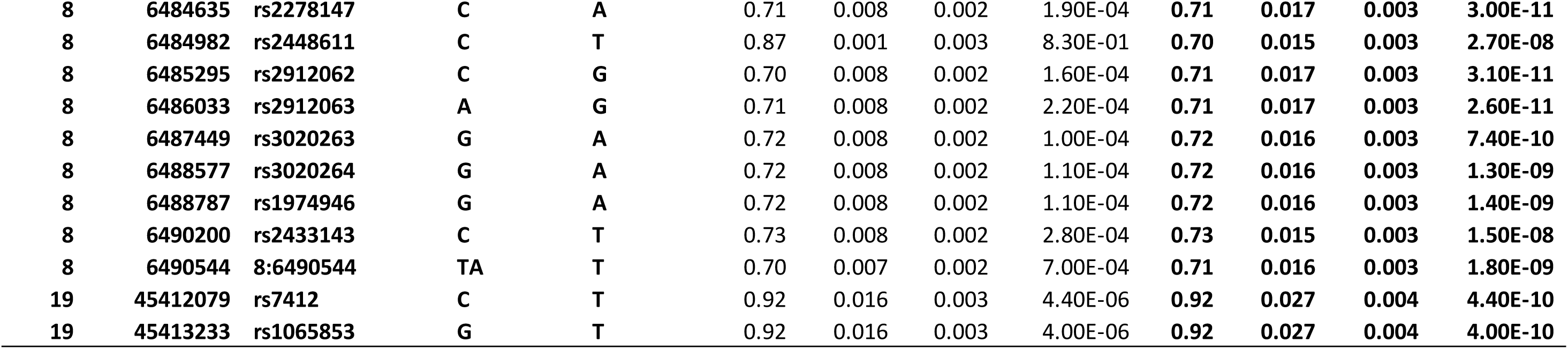
Comparison of GWAS significant SNPs for IMTmean in men and women

**Supplemental Table 5:**
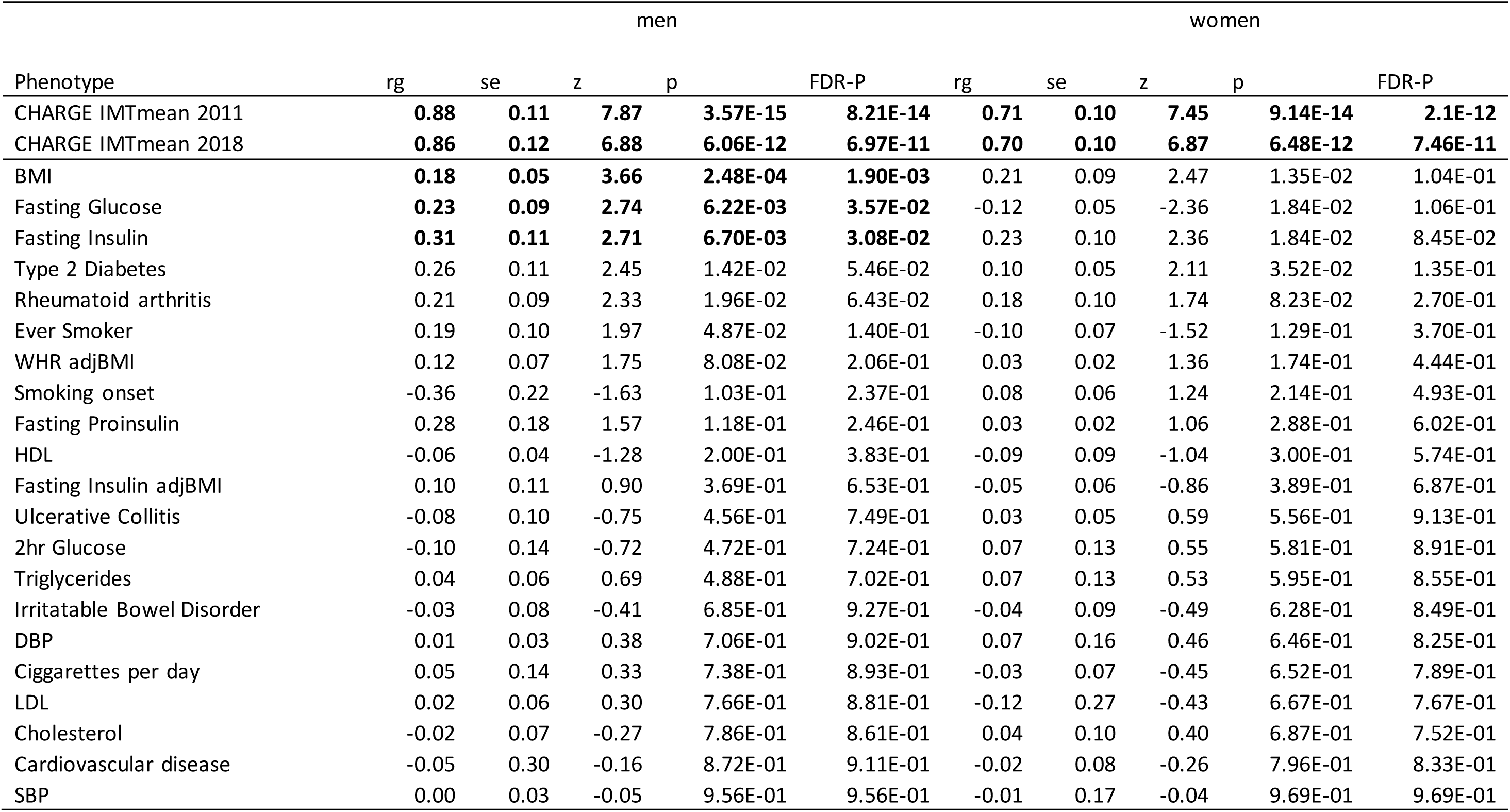
Genetic correlations with IMTmean by sex

**Supplemental Table 6:**
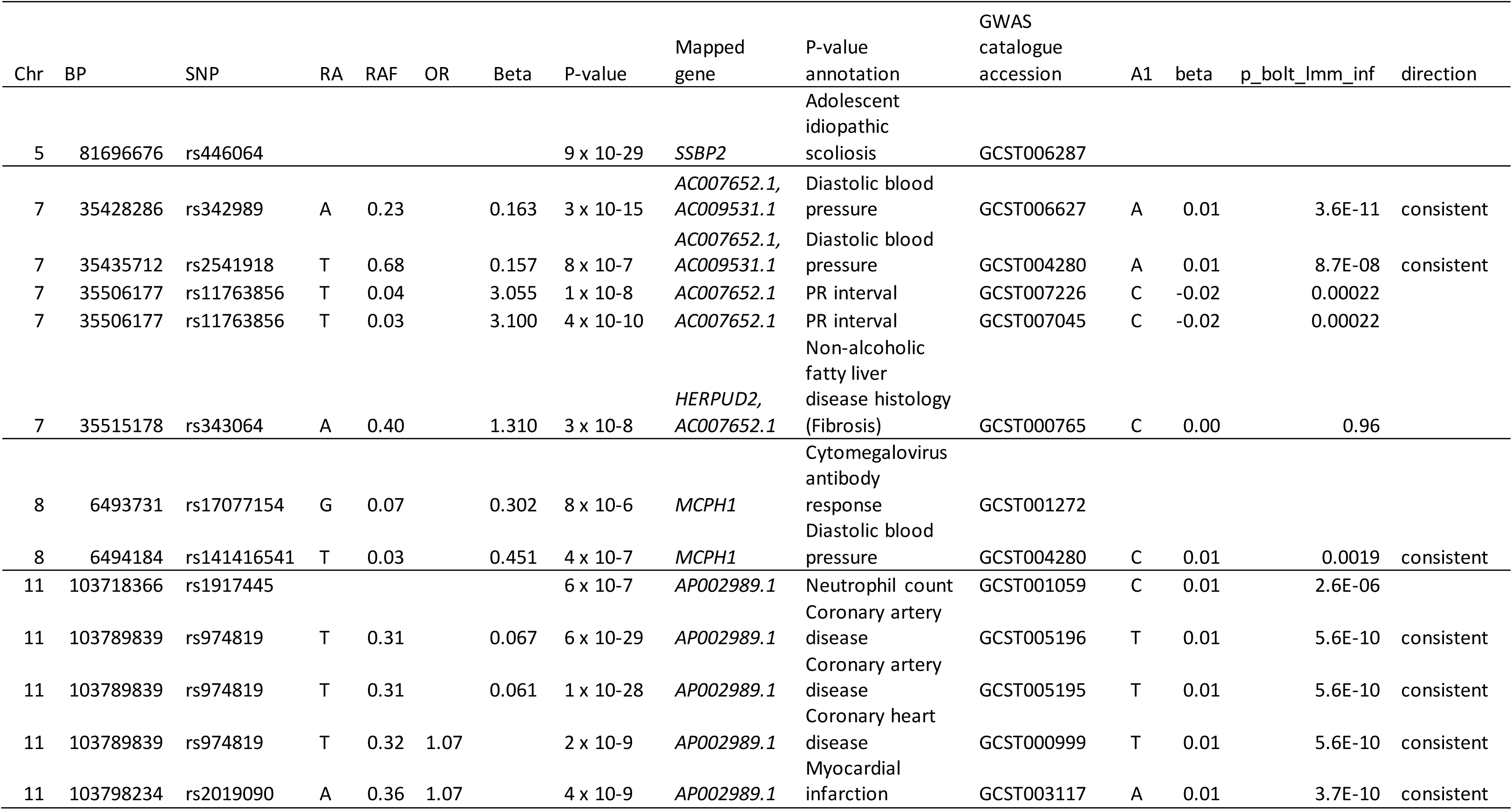
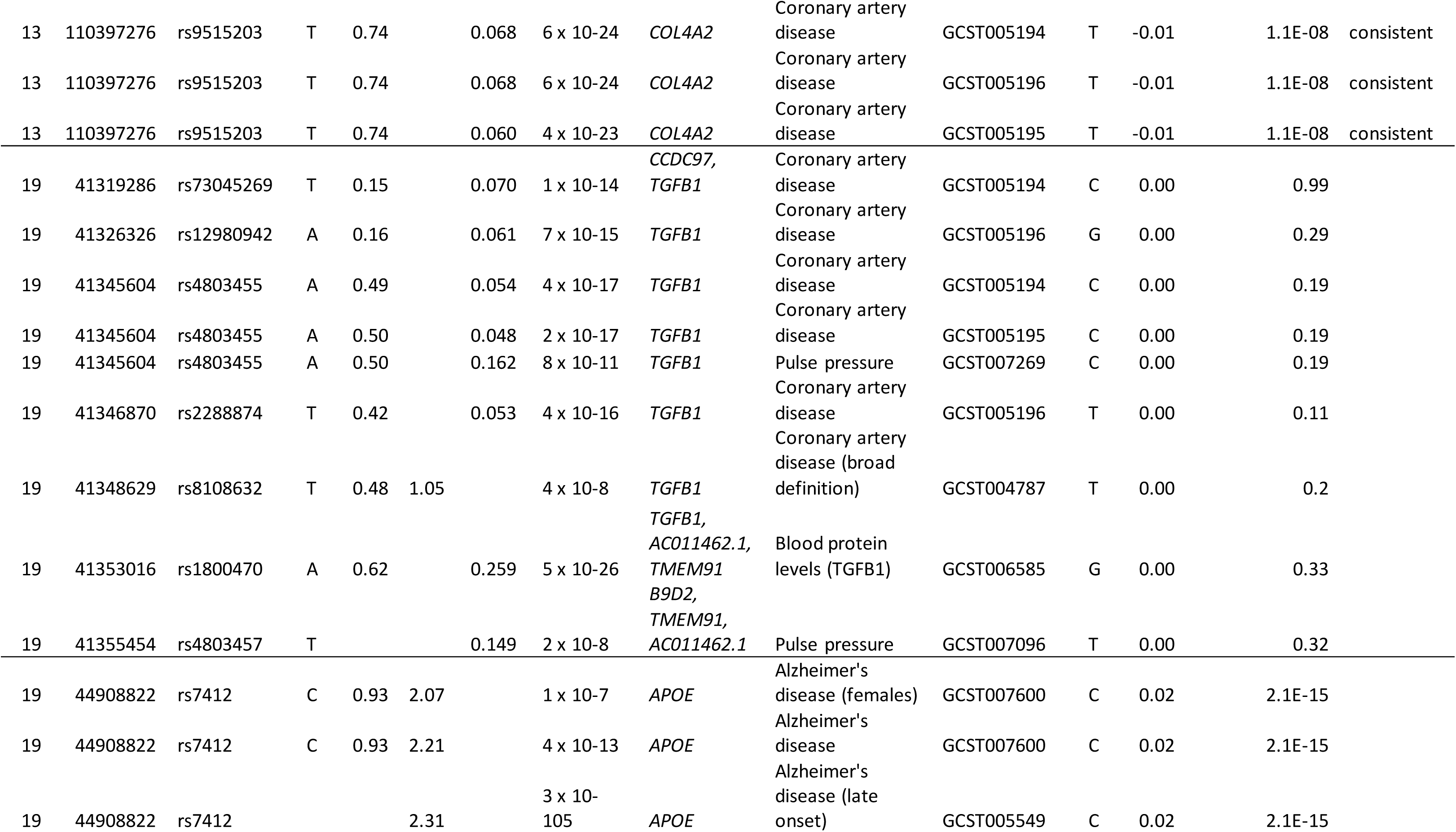
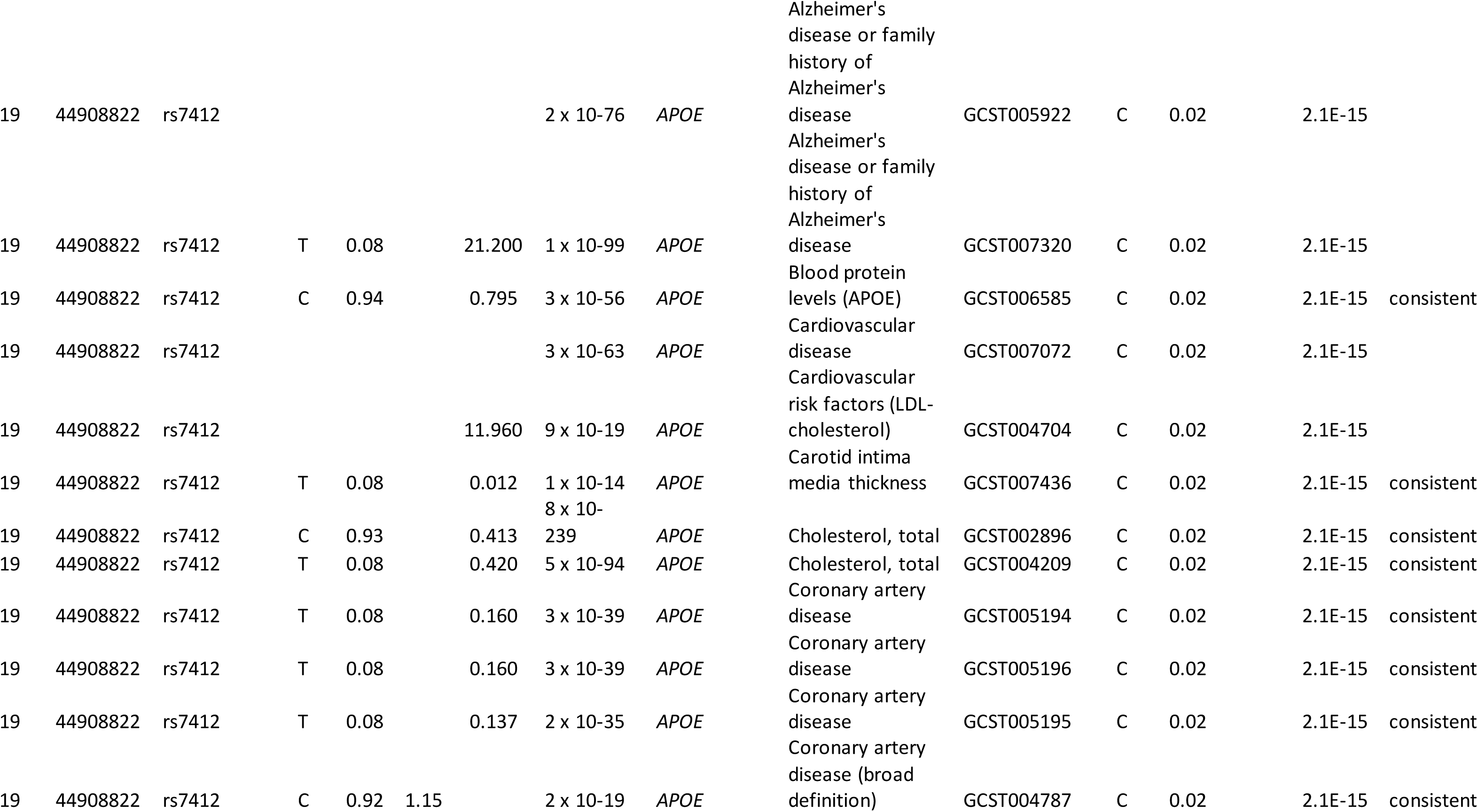
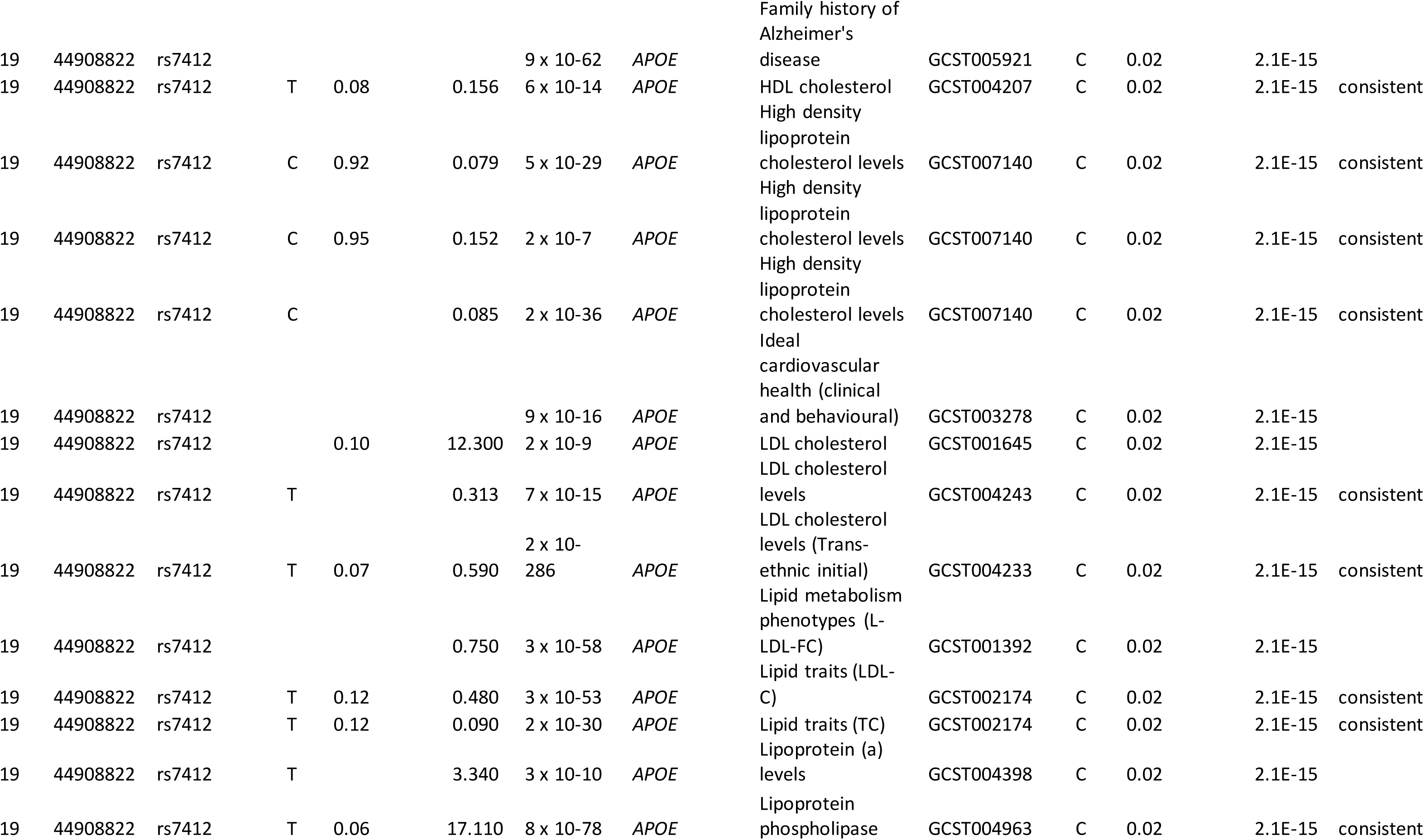
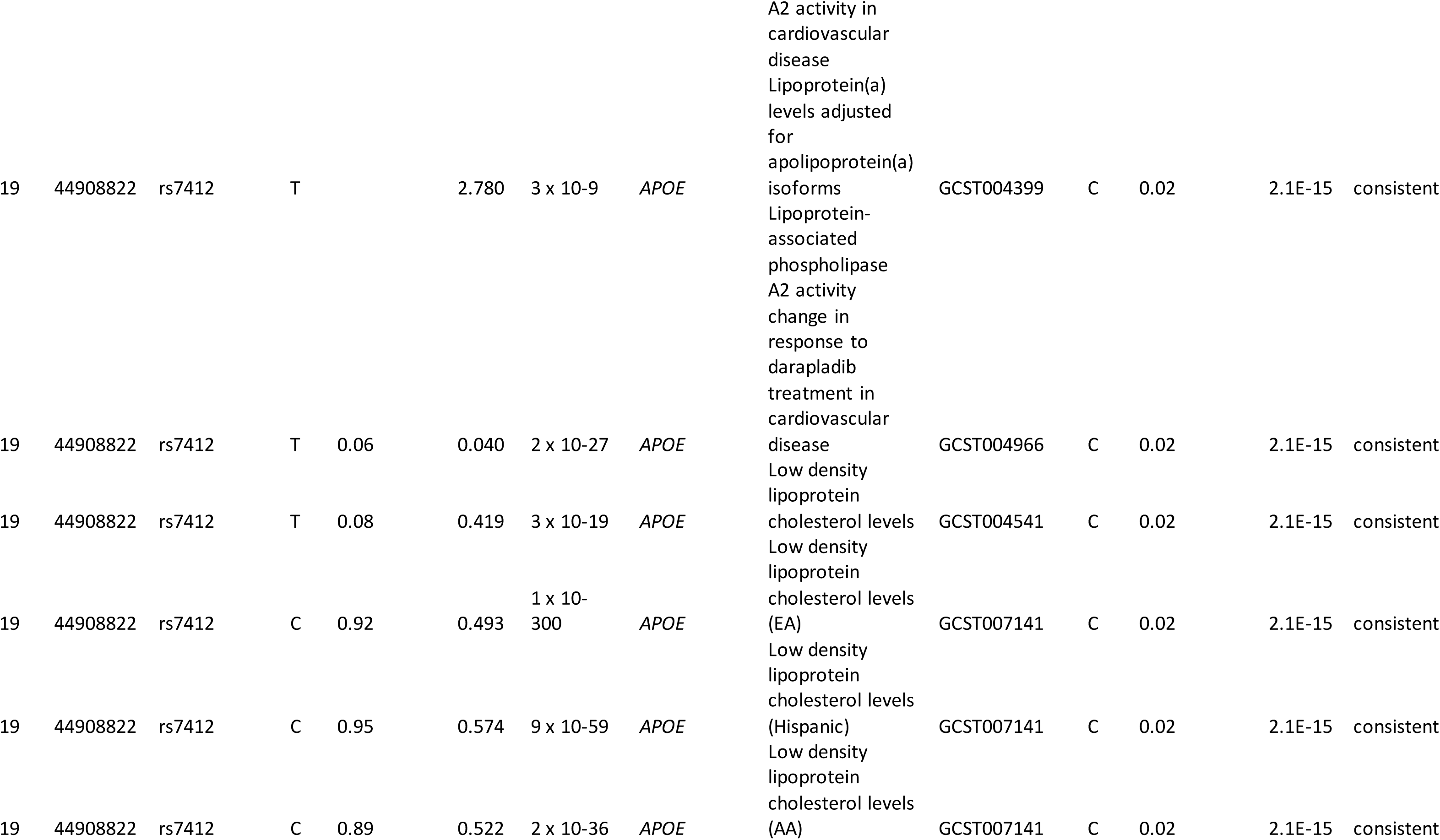
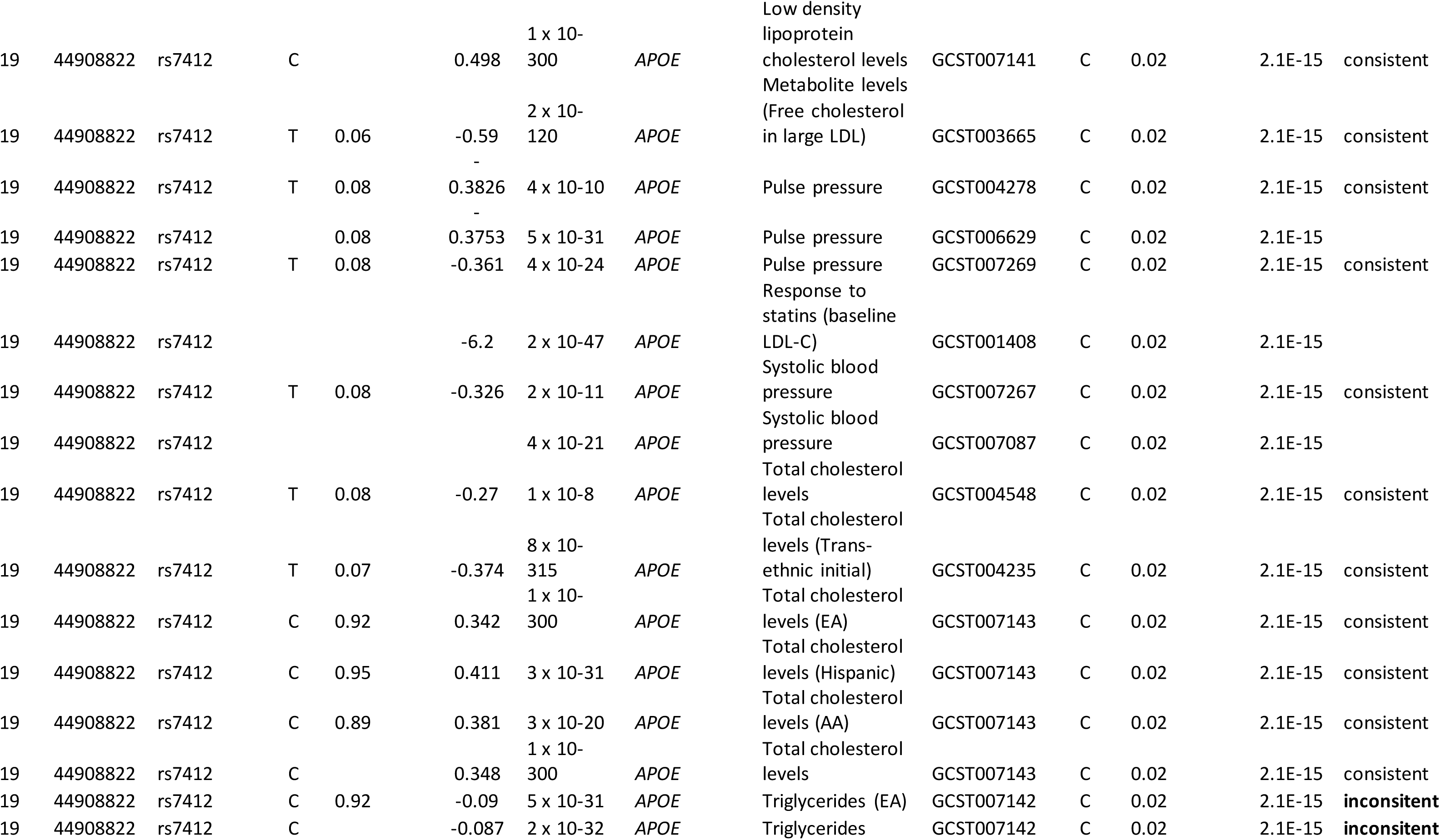
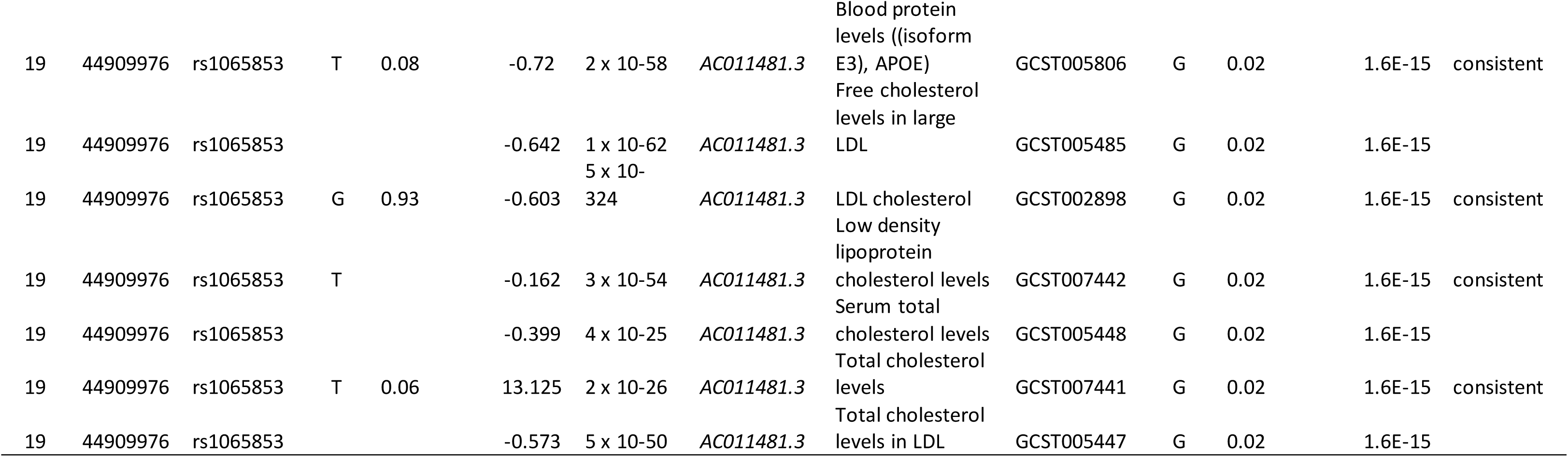
Previous associations with SNPs in GWAS-significant loci

**Supplemental Table 7:** predicted functinoal, coding or loss of function variants

| SNP | chr | pos | Allele | Consequence | IMPACT | SYMBOL | SIFT | PolyPhen |
| --- | --- | --- | --- | --- | --- | --- | --- | --- |
| rs6658335 | 1 | 1707555 | T | missense_variant | MODERATE | *CDK11A* | tolerated_low_confidence(0.06) | benign(0.007) |
| rs2912010 | 8 | 6621465 | T | synonymous_variant | LOW | *MCPH1* | - | - |
| rs1057090 | 8 | 6621521 | A | missense_variant | MODERATE | *MCPH1* | tolerated(0.08) | benign(0.129) |
| rs1057090 | 8 | 6621521 | G | missense_variant | MODERATE | *MCPH1* | deleterious(0.02) | benign(0.025) |
| rs1057090 | 8 | 6621521 | T | missense_variant | MODERATE | *MCPH1* | tolerated(1) | benign(0) |
| rs2912016 | 8 | 6621657 | A | synonymous_variant | LOW | *MCPH1* | - | - |
| rs2912057 | 8 | 6625961 | G | missense_variant | MODERATE | *MCPH1* | - | - |
| rs2912057 | 8 | 6625961 | G | missense_variant | MODERATE | *MCPH1* | - | - |
| rs11558719 | 10 | 72213199 | T | missense_variant | MODERATE | *ASCC1* | tolerated(0.21) | benign(0.203) |
| rs28375406 | 16 | 88930433 | G | missense_variant | MODERATE | *LOC105371409* | - | - |
| rs12924185 | 16 | 88950476 | T | missense_variant | MODERATE | *LOC100129697* | tolerated_low_confidence(0.07) | unknown(0) |
| rs1160983 | 19 | 44893972 | A | synonymous_variant | LOW | *TOMM40* | - | - |
| rs7412 | 19 | 44908822 | T | missense_variant | MODERATE | *APOE* | deleterious(0) | probably_damaging(1) |

**Supplemental Table 8:** EQTLS identified in GTEx

|  | SNP Id | Chr | Pos | A1 | A2 | Gene Symbol | Gencode Id | P-Value | NES | Tissue |
| --- | --- | --- | --- | --- | --- | --- | --- | --- | --- | --- |
| lead | rs309563 | 5 | 82896644 | A | T | *VCAN-AS1* | ENSG00000249835.2 | 4.60E-06 | -0.2 | Testis |
| VEP | rs1057090 | 8 | 6479042 | C | T | *MCPH1-AS1* | ENSG00000271743.1 | 5.20E-06 | -0.44 | Testis |
| VEP | rs1057090 | 8 | 6479042 | C | T | *MCPH1-AS1* | ENSG00000271743.1 | 2.70E-05 | -0.26 | Thyroid |
| VEP | rs2912010 | 8 | 6478986 | C | T | *MCPH1-AS1* | ENSG00000271743.1 | 8.30E-06 | -0.28 | Nerve - Tibial |
| VEP | rs2912010 | 8 | 6478986 | C | T | *MCPH1-AS1* | ENSG00000271743.1 | 8.00E-06 | -0.43 | Testis |
| VEP | rs2912016 | 8 | 6479178 | C | A | *MCPH1-AS1* | ENSG00000271743.1 | 3.90E-06 | -0.37 | Colon - Transverse |
| VEP | rs2912016 | 8 | 6479178 | C | A | *MCPH1-AS1* | ENSG00000271743.1 | 1.30E-05 | -0.3 | Esophagus - Muscularis |
| VEP | rs2912016 | 8 | 6479178 | C | A | *MCPH1-AS1* | ENSG00000271743.1 | 4.30E-09 | -0.37 | Nerve - Tibial |
| VEP | rs2912016 | 8 | 6479178 | C | A | *MCPH1-AS1* | ENSG00000271743.1 | 3.10E-06 | -0.58 | Prostate |
| VEP | rs2912016 | 8 | 6479178 | C | A | *MCPH1-AS1* | ENSG00000271743.1 | 3.90E-06 | -0.46 | Stomach |
| VEP | rs2912016 | 8 | 6479178 | C | A | *MCPH1-AS1* | ENSG00000271743.1 | 2.80E-06 | -0.47 | Testis |
| VEP | rs2912016 | 8 | 6479178 | C | A | *MCPH1-AS1* | ENSG00000271743.1 | 1.00E-07 | -0.33 | Thyroid |
| VEP | rs2912057 | 8 | 6483482 | C | T | *MCPH1-AS1* | ENSG00000271743.1 | 2.30E-05 | -0.26 | Nerve - Tibial |
| lead | rs2912062 | 8 | 6485295 | C | G | *MCPH1-AS1* | ENSG00000271743.1 | 3.10E-06 | -0.52 | Brain - Cortex |
| lead | rs2912062 | 8 | 6485295 | C | G | *MCPH1-AS1* | ENSG00000271743.1 | 7.70E-07 | -0.46 | Brain - Nucleus<br>accumbens (basal<br>ganglia) |
| lead | rs2912062 | 8 | 6485295 | C | G | *MCPH1-AS1* | ENSG00000271743.1 | 1.20E-06 | -0.65 | Cells - EBV-transformed<br>lymphocytes |
| lead | rs2912062 | 8 | 6485295 | C | G | *MCPH1-AS1* | ENSG00000271743.1 | 5.40E-11 | -0.55 | Colon - Transverse |
| lead | rs2912062 | 8 | 6485295 | C | G | *MCPH1-AS1* | ENSG00000271743.1 | 1.60E-06 | -0.35 | Esophagus - Muscularis |
| lead | rs2912062 | 8 | 6485295 | C | G | *MCPH1-AS1* | ENSG00000271743.1 | 8.70E-11 | -0.43 | Nerve - Tibial |
| lead | rs2912062 | 8 | 6485295 | C | G | *MCPH1-AS1* | ENSG00000271743.1 | 2.70E-06 | -0.58 | Ovary |
| lead | rs2912062 | 8 | 6485295 | C | G | *MCPH1-AS1* | ENSG00000271743.1 | 2.10E-06 | -0.48 | Stomach |
| lead | rs2912062 | 8 | 6485295 | C | G | *MCPH1-AS1* | ENSG00000271743.1 | 4.80E-08 | -0.55 | Testis |
| lead | rs2912062 | 8 | 6485295 | C | G | *MCPH1-AS1* | ENSG00000271743.1 | 2.40E-07 | -0.34 | Thyroid |
| lead | rs2912063 | 8 | 6486033 | A | G | *MCPH1-AS1* | ENSG00000271743.1 | 4.60E-06 | -0.51 | Brain - Cortex |
| lead | rs2912063 | 8 | 6486033 | A | G | *MCPH1-AS1* | ENSG00000271743.1 | 2.00E-06 | -0.44 | Brain - Nucleus<br>accumbens (basal<br>ganglia) |
| lead | rs2912063 | 8 | 6486033 | A | G | *MCPH1-AS1* | ENSG00000271743.1 | 8.40E-11 | -0.55 | Colon - Transverse |
| lead | rs2912063 | 8 | 6486033 | A | G | *MCPH1-AS1* | ENSG00000271743.1 | 3.80E-07 | -0.37 | Esophagus - Muscularis |
| lead | rs2912063 | 8 | 6486033 | A | G | *MCPH1-AS1* | ENSG00000271743.1 | 6.20E-10 | -0.41 | Nerve - Tibial |
| lead | rs2912063 | 8 | 6486033 | A | G | *MCPH1-AS1* | ENSG00000271743.1 | 2.90E-06 | -0.59 | Ovary |
| lead | rs2912063 | 8 | 6486033 | A | G | *MCPH1-AS1* | ENSG00000271743.1 | 1.20E-06 | -0.5 | Stomach |
| lead | rs2912063 | 8 | 6486033 | A | G | *MCPH1-AS1* | ENSG00000271743.1 | 1.10E-07 | -0.54 | Testis |
| lead | rs2912063 | 8 | 6486033 | A | G | *MCPH1-AS1* | ENSG00000271743.1 | 3.30E-07 | -0.34 | Thyroid |
| lead | rs2019090 | 11 | 103668962 | A | T | *RP11-563P16.1* | ENSG00000254987.1 | 3.70E-05 | -0.3 | Artery - Aorta |
| lead | rs561732 | 16 | 88988989 | T | C | *CBFA2T3* | ENSG00000129993.10 | 7.10E-15 | -0.47 | Pancreas |
| lead | rs561732 | 16 | 88988989 | T | C | *CBFA2T3* | ENSG00000129993.10 | 1.90E-05 | -0.17 | Thyroid |
| VEP | rs28375406 | 16 | 88996841 | A | G | *CBFA2T3* | ENSG00000129993.10 | 2.90E-10 | -0.39 | Pancreas |
| VEP | rs28375406 | 16 | 88996841 | A | G | *CBFA2T3* | ENSG00000129993.10 | 1.50E-05 | -0.17 | Thyroid |
| VEP | rs7412 | 19 | 45412079 | C | T | *APOE* | ENSG00000130203.5 | 2.20E-05 | -0.29 | Skin - Not Sun Exposed<br>(Suprapubic) |

Where: CTD-2541M15.3 also known as MCPH1-AS1

